# PCNA encircling primer/template junctions is eliminated by exchange of RPA for Rad51: Implications for the interplay between human DNA damage tolerance pathways

**DOI:** 10.1101/2025.03.27.645792

**Authors:** Jessica L. Norris, Lindsey O. Rogers, Grace Young, Kara G. Pytko, Rachel L. Dannenberg, Samuel Perreault, Vikas Kaushik, Edwin Antony, Mark Hedglin

## Abstract

The DNA genome is constantly exposed to agents, such as ultraviolet radiation (UVR), that can alter or eliminate its coding properties through covalent modifications of the template bases. Many of these damaging modifications (i.e., lesions) persist into S-phase of the cell cycle where they may stall the canonical DNA replication machinery. In humans, these stalling events are circumvented by one of at least three interconnected DNA damage tolerance (DDT) pathways; translesion DNA synthesis (TLS), Template Switching (TS), and Homology-dependent Recombination (HDR). Currently, the functional interplay between these pathways is unclear, leaving wide gaps in our fundamental understanding of human DDT. To gain insights, we focus on the activation mechanisms of the DDT pathways. PCNA sliding clamps encircling primer/template (P/T) junctions of stalled replication sites are central to the activation of both TLS and TS whereas exchange of RPA for Rad51 filaments on the single strand DNA (ssDNA) sequences of stalled replication sites is central to HDR activation. Utilizing direct, ensemble FRET approaches developed by our lab, we independently monitor and directly compare PCNA occupancy and RPA/Rad51 exchange at P/T junctions under a variety of conditions that mimic *in vivo* scenarios. Collectively, the results reveal that assembly of stable Rad51 filaments at P/T junctions via RPA/Rad51 exchange causes complete and irreversible unloading of the resident PCNA, both in the presence and absence of an abundant PCNA-binding protein complex. Further investigations decipher the mechanism of RPA/Rad51 exchange-dependent unloading of PCNA. Collectively, these studies provide critical insights into the interplay between human DDT pathways and direction for future studies.

## Introduction

Proper genetic inheritance requires that the double strand DNA (dsDNA) genome of each cell is accurately replicated for transfer to its progeny. The bulk of DNA replication is carried out by a canonical pathway that relies on the high-fidelity DNA polymerases (pols), ε and δ, and essential accessory factors such as the processivity sliding clamp ring, proliferating cell nuclear antigen (PCNA), the PCNA clamp loader, replication factor C (RFC), and the major single strand DNA (ssDNA) binding protein complex, replication protein A (RPA) (**Figure 1A**). RFC utilizes ATP to load PCNA onto primer/template (P/T) junctions such that the “front face” of loaded PCNA is oriented towards the 3′ terminus of the primer strand from where DNA synthesis initiates from (i.e., in the direction of DNA synthesis)^1–4^. Pols ε and δ are responsible for the replication of leading and lagging strand ssDNA templates, respectively, and like many DNA processing enzymes, anchor to the front face of loaded PCNA forming holoenzymes with maximal efficiency. RPA directly engages P/T junctions and coats ssDNA templates as they are exposed during replication, forming directional filaments that protect the underlying ssDNA from being degraded or forming secondary structures that limit its accessibility^5–12^. The affinity of human RPA for dsDNA is nearly 3 orders of magnitude weaker than its affinity for ssDNA ^13–15^. Hence, conversion of template ssDNA to dsDNA by DNA pols indirectly and rapidly releases RPA into solution such that RPA filaments are short and transient under non-stressed conditions^15–18^.

**Figure 1.**
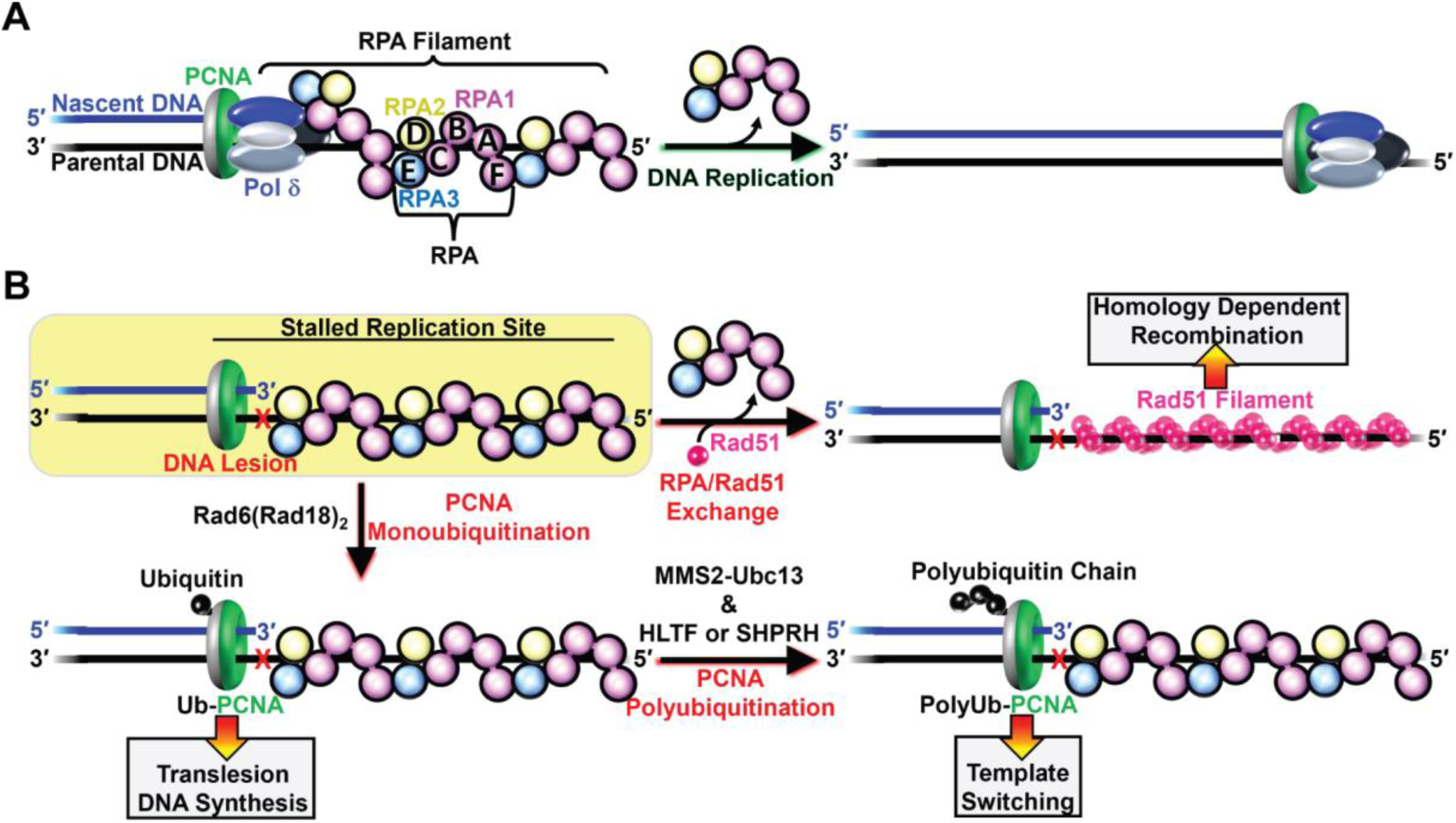
DNA damage and human DNA replication. Nascent DNA (primer) and parental DNA (template) strands are blue and black, respectively. RPA subunits are color-coded and depicted to illustrate the OB-folds (A – E). The “front” and “back” faces of PCNA are green and grey, respectively. For clarity, RFC is not shown. **A**) Canonical DNA replication. Shown as an example is replication of lagging strand DNA template by pol δ. The ssDNA template to be replicated is extended by an RPA filament in which each RPA engages the ssDNA in an orientation-specific manner (as depicted). **B**) DNA damage tolerance. All pathways emanate from a site of stalled canonical DNA replication (highlighted in yellow). The offending DNA lesion is indicated by an “X.” Activation of a given pathway is indicated by →.

The DNA genome is constantly exposed to agents, such as ultraviolet radiation (UVR), that can alter or eliminate its coding properties through covalent modifications of the template bases. Many of these damaging modifications (i.e., DNA lesions), such as the major photoproducts generated from UVR exposure, cannot be accommodated by pols ε and δ and, consequently, stall canonical replication, generating long, persistent RPA filaments that directly engage the afflicted P/T junctions and the unreplicated ssDNA templates downstream. The resident PCNA are left behind on the dsDNA regions upstream of the afflicted P/T junctions ^19,20^. In humans, these stalling events are circumvented by one of at least three non-canonical DNA replication pathways that bypass (i.e., “tolerate”) the offending lesion, permitting canonical DNA replication to resume downstream^21–23^. This process is referred to as DNA damage tolerance (DDT) and is imperative for cell survival following mild exposure to DNA damaging agents. Previous studies have identified the fundamental components of each DDT pathway and deciphered their basic mechanisms and interdependencies (**Figure 1B**)^21–25^.

Translesion DNA synthesis (TLS) relies on the attachment of single ubiquitin moieties to K164 residue(s) on the “back face” of PCNA encircling sites of stalled canonical DNA replication (i.e., stalled replication site). This posttranslational modification (PTM), referred to as PCNA monoubiquitination, is catalyzed by the Rad6(Rad18)_2_ complex and activates TLS by recruiting essential TLS factors to stalled replication sites. TLS culminates with one or more specialized DNA pols (i.e., TLS pols) utilizing monoubiquitinated PCNA to extend the aborted primer strand directly opposite and beyond the offending lesion. TLS pols all contain expanded active sites that accommodate an array of DNA lesions, albeit with lowered fidelities^23,24,26,27^. In template switching (TS), the single ubiquitin moieties attached to the “back face” of PCNA by the Rad6(Rad18)_2_ complex are extended to a K63-linked polyubiquitin chain by the MMS2-Ubc13 complex in conjunction with either HLTF or SHPRH. This PTM, referred to as PCNA polyubiquitination, activates TS by recruiting the AH2/ZRANB3 DNA translocase, which subsequently catalyzes replication fork reversal, displacing RPA. The resultant Holliday junction-like structure permits an unidentified DNA pol to utilize the nascent DNA strand from the undamaged, homologous sister chromatid as a template to extend the aborted primer^28–31^. In homology-dependent recombination (HDR), RPA is exchanged for ATP-bound Rad51 (i.e., Rad51•ATP) on the unreplicated ssDNA templates of stalled replication sites. Rad51 is an ATP-dependent DNA recombinase and activates HDR by catalyzing strand invasion in which the aborted primer strand invades the undamaged, homologous sister chromatid at a complimentary sequence. In this context, Rad51-catalyzed strand invasion results in a multi-branched, H-shaped structure. An unidentified DNA pol then utilizes the nascent DNA strand from the undamaged, homologous sister chromatid as a template to extend the aborted primer. Unlike TLS and TS, HDR is independent of PCNA monoubiquitination and a recent study suggests PCNA does not serve a functional role in Rad51-catalyzed strand invasion during HDR^19,21,22,32^. In both TS and HDR pathways, the aborted primer strands are extended to a degree such that when they are subsequently re-annealed to the damaged template strand (via reversal of the higher-order DNA structures) their 3′ termini are placed downstream of the offending lesion.

All DDT pathways emanate from stalled replication sites, contribute to cell survival following exposure to common DNA damaging agents such as UVR, and, if successfully executed, yield the same results. Namely, faithful replication of a DNA lesion and the contiguous DNA sequence and the resumption of canonical DNA replication downstream of the offending lesion. Currently, the functional interplay between DDT pathways at stalled replication sites is unclear, leaving wide gaps in our fundamental understanding of human DDT. To gain insights we focus on the basic activation mechanisms of DDT pathways. PCNA encircling P/T junctions of stalled replication sites is central to the activation of both TLS and TS, as described above and depicted in **Figure 1B**. Multiple mechanisms collectively maintain loaded PCNA at/near these sites. For example, “protein roadblocks” restrict diffusion of loaded PCNA away from stalled replication sites^15,33^. Large DNA-binding protein complexes that rapidly and tightly engage nascent DNA as it is generated during DNA replication restrict diffusion of the resident PCNA along the dsDNA upstream of stalled replication sites^34–36^. RPA•ssDNA complexes immediately adjacent to P/T junctions of stalled replication sites prevent diffusion of loaded PCNA onto and along the unreplicated ssDNA templates^4,37^. Furthermore, these RPA•ssDNA complexes prohibit RFC-catalyzed unloading of PCNA and permit RFC-catalyzed re-loading of PCNA back onto stalled replication sites that are devoid of the sliding clamp^4,15,24^. Exchange of RPA for Rad51•ATP filaments on the unreplicated ssDNA templates of stalled replication sites is central to HDR activation (**Figure 1B**). In the present study, we utilize direct, ensemble FRET approaches developed by our lab to investigate the interplay between RPA/Rad51 exchange and PCNA occupancy at P/T junctions under a variety of conditions that mimic *in vivo* scenarios^15,38^. Collectively, the results from the present study provide critical insights into the functional interplay between human DDT pathways and direction for future studies.

## Materials and Methods

### Oligonucleotides

Oligonucleotides were synthesized by Integrated DNA Technologies (Coralville, IA) and purified via denaturing polyacrylamide gel electrophoresis. The concentrations of unlabeled DNAs were determined from the absorbance at 260 nm using the provided extinction coefficients. The concentrations of Cy5-labeled DNAs were determined from the extinction coefficient at 650 nm for Cy5 (ε_650_ = 250,000 M^−1^cm^−1^). Concentrations of Cy3-labeled DNAs were determined from the extinction coefficient at 550 nm for Cy3 (ε_550_ = 136,000 M^−1^cm^−1^). For annealing two ssDNAs (as depicted in **Figure S1**), the primer and corresponding complementary template strands were mixed in equimolar amounts in 1X Annealing Buffer (10 mM TrisHCl, pH 8.0, 100 mM NaCl, 1 mM EDTA), heated to 95 °C for 5 minutes, and allowed to slowly cool to room temperature.

### Recombinant Human Proteins

Human RPA, Cy5-PCNA, exonuclease-deficient Pol δ (referred to herein as simply Pol δ) and RFC were obtained as previously described^2,39–41^. The plasmid (pCH1/RAD51o) for the expression of wild-type human Rad51 was a gift from Maria Spies (Addgene plasmid #102562; http://n2t.net/addgene:102562; RRID:Addgene_102562) and human Rad51 was obtained as previously described^41^. Human RPA containing a Cy5 label at residue 215 of the oligonucleotide binding (OB) fold A of the RPA1 subunit (RPA-OBA-Cy5) was obtained as described previously^15,42^. The concentrations of active RPA, RPA-OBA-Cy5, and Rad51 were determined via a FRET-based activity assays as described previously^15,41,43^.

### FRET Measurements

All FRET experiments were performed at room temperature (23 ± 2 °C) in 1X Mg^2+^/Ca^2+^ buffer (20 mM HEPES, pH 7.5, 150 mM KCl, 5 mM MgCl_2_, 5 mM CaCl_2_) supplemented with 1 mM DTT, 1 mM ATP, and the ionic strength was adjusted to physiological (200 mM) by the addition of appropriate amounts of KCl. Ca^2+^ is included to stabilize active Rad51 nucleoprotein filaments on ssDNA via restriction of ATP hydrolysis (but not ATP binding) by Rad51^41^. Ca^2+^ promotes more regular and stable Rad51 nucleoprotein filaments on ssDNA^41,44^ but does not affect the amount of RPA that binds to ssDNA nor the amount of PCNA loaded onto DNA by RFC via a reaction mechanism that requires ATP binding and hydrolysis^15^. All experiments were performed in a 16.100F-Q-10/Z15 sub-micro fluorometer cell (Starna Cells) and monitored in a Horiba Scientific Duetta-Bio fluorescence/absorbance spectrometer as described previously^15^. In short, reaction solutions are excited at 514 nm and the fluorescence emission intensities (*I*) are monitored essentially simultaneously (Δt = 0.118 ms) at the peak emission wavelengths for Cy3 (563 nm, *I*_563_) and Cy5 (665 nm, *I*_665_) over time, recording *I* every 0.17 s. For all experiments, excitation and emission slit widths are 10 nm. All recorded fluorescence emission intensities are corrected by a respective dilution factor and all time courses are adjusted for the time between the addition of each component and the fluorescence emission intensity recordings (i.e., dead time ≤ 10 s). For any recording of the fluorescence emission intensities (*I_665_* and *I_563_*), the approximate FRET efficiency is estimated from the equation [inline]. For each experiment below, the final concentrations of all reaction components are indicated. The concentrations of ATP in all experimental solutions described below is 1.0 mM and, hence, this concentration is maintained upon mixing.

### Pre-steady state FRET experiments to monitor RPA/Rad51 exchange

For experiments performed in the absence of PCNA, RFC, and pol δ, a solution containing ddP/5′TCy3 DNA (20 nM, **Figure S1**), NeutrAvidin (80 nM homotetramer), and ATP is pre-incubated with an RPA (25 nM active heterotrimer, RPA or RPA-OBA-Cy5), the resultant solution is transferred to a fluorometer cell, and the cell is placed in the instrument. *I*_665_ and *I_563_* are monitored over time until both stabilize for at least 1 min. Within this stable region, E_FRET_ values are calculated from the observed *I*_665_ and *I_563_* values and averaged to obtain the E_FRET_ value observed prior to addition of Rad51•ATP. Then, pre-formed Rad51•ATP (500 nM active Rad51 monomer, ATP) is added, the resultant solution is mixed by pipetting, and *I*_665_ and *I_563_* are monitored beginning ≤ 10 s after the addition of Rad51•ATP.

Experiments in the presence of PCNA and RFC are performed as described above except that PCNA is first pre-loaded onto DNA, as previously described^15^. In short, an RPA (25 nM active heterotrimer, RPA or RPA-OBA-Cy5) is added to a solution containing ddP/5′TCy3 DNA (20 nM, **Figure S1**), NeutrAvidin (80 nM homotetramer), and ATP. PCNA (20 nM homotrimer, PCNA or Cy5-PCNA) and ATP-bound RFC (i.e., RFC•ATP, 20 nM RFC heteropentamer, ATP) are then added in succession. RPA-OBA-Cy5 fully supports RFC-catalyzed loading of PCNA onto P/T junctions^15^. The resultant solution is transferred to a fluorometer cell, and the cell is placed in the instrument. 5 min after the addition of RFC•ATP (i.e., PCNA loading completed), *I*_665_ and *I_563_* are monitored over time until both stabilize for at least 1 min. Within this stable region, E_FRET_ values are calculated from the observed *I*_665_ and *I_563_* values and averaged to obtain the E_FRET_ value observed prior to addition of Rad51•ATP. Finally, pre-formed Rad51•ATP (500 nM active Rad51 monomer, ATP) is added and the experiment is continued as described above.

Experiments in the presence of PCNA, RFC, pol δ, +/− dGTP experiments are performed as described above except pol δ holoenzymes are first pre-assembled on P/T DNA, as previously described^15^. In short, a PCNA (20 nM homotrimer, PCNA or Cy5-PCNA) is loaded by RFC•ATP (20 nM RFC heteropentamer, ATP) onto ddP/5′TCy3 DNA (20 nM, **Figure S1**) that is pre-engaged by NeutrAvidin (80 nM homotetramer) and an RPA (25 nM active heterotrimer, RPA or RPA-OBA-Cy5) exactly as described above. 5 min after the addition of RFC•ATP (i.e., PCNA loading complete), dGTP (0 or 1.0 mM) and pol δ (40 nM heterotetramer) are added in immediate succession, the resultant solution is transferred to a fluorometer cell, and the cell is placed in the instrument. 5 min after the addition of pol δ +/− dGTP (i.e., formation of pol δ holoenzymes or adoption of initiation states of pol δ holoenzymes for DNA synthesis is completed), *I*_665_ and *I_563_* are monitored over time until both stabilize for at least 1 min. Within this stable region, E_FRET_ values are calculated from the observed *I*_665_ and *I_563_* values and averaged to obtain the E_FRET_ value observed prior to addition of Rad51•ATP. Finally, pre-formed Rad51•ATP (500 nM active Rad51 monomer, ATP) are added and the experiment is continued as described above.

To determine the predicted E_FRET_ trace for RPA-OBA-Cy5 remaining completely disengaged from the ddP/5′TCy3 DNA substrate in the aforementioned experiments, *I*_665_ and *I*_563_ for RPA-OBA-Cy5 alone (*I*^*RPA*^_665_ and *I*^*RPA*^_563_) and a corresponding reaction mixture lacking only RPA-OBA-Cy5 (*I*^*DNA*^_665_ and *I*^*DNA*^_563_) are each monitored, and E_FRET_ is calculated as follows:

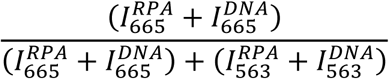

### Pre-steady FRET Experiments to monitor PCNA Loading

Experiments in the presence of RPA are carried out exactly as described in a previous report^15^. In short, RPA (25 nM active heterotrimer) is added to a solution containing 5′ddPCy3/T DNA (20 nM, **Figure S1**), NeutrAvidin (80 nM homotetramer), and ATP. Cy5-PCNA (20 nM homotrimer) is added, the resultant solution is transferred to a fluorometer cell, and the cell is placed in the instrument. *I*_665_ and *I_563_* are monitored over time until both stabilize for at least 1 min. Within this stable region, E_FRET_ values are calculated from the observed *I*_665_ and *I_563_* values and averaged to obtain the E_FRET_ value observed prior to addition of RFC•ATP. Finally, RFC•ATP (20 nM RFC heteropentamer, ATP) is added, the resultant solution is mixed by pipetting, and *I*_665_ and *I_563_* are monitored beginning ≤ 10 s after the addition of RFC•ATP. Experiments in the presence of Rad51•ATP are carried out exactly as described above except that RPA is replaced with pre-formed Rad51•ATP (200 nM active Rad51 monomer, ATP).

To determine the predicted E_FRET_ trace for Cy5-PCNA remaining completely disengaged from 5′ddPCy3/T nucleoprotein complexes/filaments (i.e., Rad51 and RPA present) in the aforementioned experiments, *I*_665_ and *I*_563_ for Cy5-PCNA alone (*I*^*PCNA*^_665_ and *I*^*PCNA*^_665_) and a corresponding reaction mixture lacking only Cy5-PCNA (*I*^*PCNA*^_563_ and *I*^*DNA*^_563_) are each monitored, and E_FRET_ is calculated as follows:

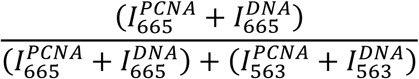

## Results

### PCNA encircling P/T junctions engaged by RPA are irreversibly unloaded through RPA/Rad51 exchange

In a previous study, we developed ensemble FRET assays that directly and continuously monitor interactions of RPA and PCNA with P/T junctions during pol δ holoenzyme assembly and the initiation of DNA synthesis^15,38^. In the present study, we adapt these assays to monitor interactions of RPA and PCNA with P/T junctions during RPA/Rad51 exchange under a variety of conditions that mimic *in vivo* scenarios. A high-fidelity DNA polymerase, such as pol δ, dissociates into solution upon encountering DNA lesions it cannot accommodate, leaving loaded PCNA behind on the dsDNA regions upstream of the vacated P/T junctions. Thus, we initially focus on RPA/Rad51 exchange at P/T junctions encircled by PCNA (**Figure 2A**). Here, a Cy3-labeled P/T DNA substrate (**Figure S1**, ddP/5′TCy3 or 5′ddPCy3/T) is first pre-bound with NeutrAvidin and the respective RPA (RPA or RPA-OBA-Cy5). NeutrAvidin avidly binds biotin attached to the 5′ termini of the primer strands (*K*_D_ ~ pM) and the resultant complex prohibits diffusion of loaded PCNA off the blunt duplex ends of the P/T DNA substrates, emulating the aforementioned protein roadblocks upstream of stalled replication sites^2,4,34–37^.The lengths of the template ssDNA regions (33 nucleotides, nts) of the P/T DNA substrates accommodate 1 RPA heterotrimer in the 30 nucleotide binding mode (*K*_D_ ≤ 100 pM) at the RPA:DNA ratios utilized^5–7,15,45^. As indicated above, RPA promotes PCNA occupancy of P/T junctions through multiple means. Together, the biotin•NeutrAvidin and RPA•DNA complexes maintain loaded PCNA on the dsDNA regions of the P/T DNA substrates^4,15,24^. Next, a PCNA is added (PCNA or Cy5-PCNA) followed by RFC•ATP and the resultant solution is incubated until it reaches equilibrium where a PCNA (PCNA or Cy5-PCNA) is loaded and maintained on all P/T DNA•RPA complexes^4,15^. Finally, Rad51•ATP is added. All assays in the current study are performed at physiological ionic strength (200 mM I) in the absence of ATP hydrolysis by Rad51 (see **Materials and Methods**). Under these conditions, Rad51 has an exceptionally strong, intrinsic preference for forming filaments on ssDNA (**Figure S2**) such that dsDNA binding is dramatically reduced if observed at all^46–48^. The ssDNA regions of the Cy3-labeled P/T DNA substrates (**Figure S1**, ddP/5′TCy3 or 5′ddPCy3/T) accommodate Rad51•ATP filaments comprised of 11 Rad51•ATP monomersa.

**Figure 2.**
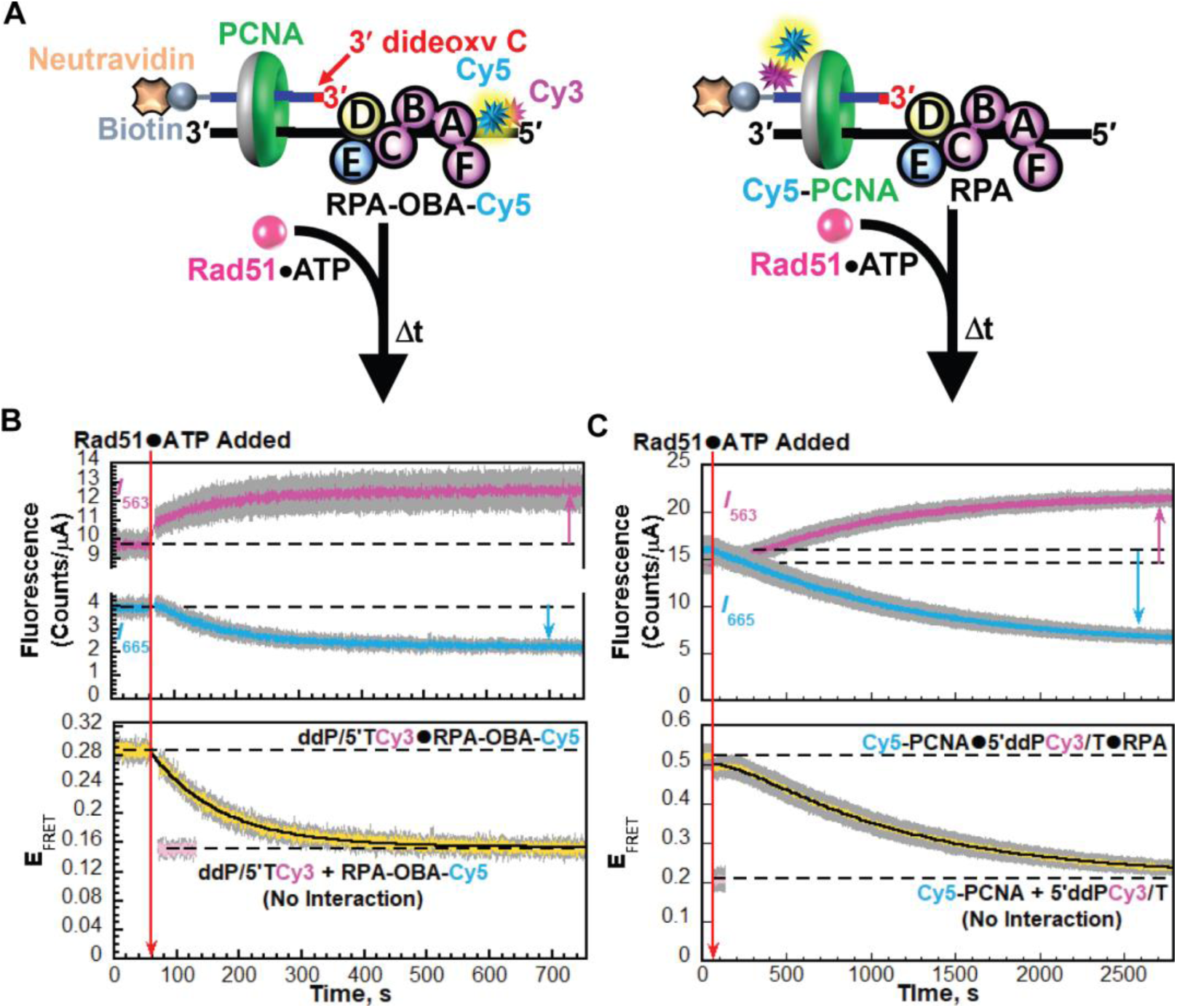
All PCNA encircling P/T junctions engaged by RPA are irreversibly unloaded through RPA/Rad51 exchange (**A**) Schematic representations of the FRET pairs and experiments to monitor the interactions of RPA (*Left*) and PCNA (*Right*) with P/T junctions during RPA/Rad51 exchange (**B - C**) Data. Each trace is the mean of at least three independent traces with the S.E.M. shown in grey. The time trajectories of *I*563 (magenta) and *I*665 (cyan) are displayed in the top panels and the corresponding EFRET (yellow) is displayed in the bottom panels. The times at which Rad51•ATP are added is indicated by red arrows. Changes in *I*563 and *I*665 are indicated in the top panels by magenta and cyan arrows, respectively. For observation, the *I*563 and *I*665, and the corresponding EFRET values observed prior to the addition of Rad51•ATP are each fit to dashed flat lines that are extrapolated to the axes limits. The predicted EFRET traces for no interaction between the respective Cyanine-labeled species are displayed in pink in each bottom panel, offset to the time after Rad51•ATP is added and fit to a dashed flat line that is extrapolated to the axis limits. Data for interactions of RPA and PCNA with P/T junctions during RPA/Rad51 exchange are shown in panels **B** and **C,** respectively. The rate constant(s) for the EFRET decrease observed after the addition of Rad51•ATP in each bottom panel is reported in **Table 1**.

For experiments monitoring interactions of RPA with P/T junctions (**Figure 2A**, *Left*), *I*_563_ and *I*_665_ values are stabilized (**Figure 2B**, *Top*) and a significant, constant E_FRET_ is observed prior to the addition of Rad51•ATP (**Figure 2B**, *Bottom*) due to the stable interaction of RPA-OBA-Cy5 with the ddP/5′TCy3 DNA^15^. Upon addition of Rad51•ATP, *I*_563_ increases simultaneously with decreases in *I*_665_ (**Figure 2B**, *Top*). These synchronized, anti-correlated changes in *I*_563_ and *I*_665_ are indicative of a direct FRET decrease due to the increase in distance between the Cy3 FRET donor on the ddP/5′TCy3 DNA substrate and the Cy5 FRET acceptor on RPA-OBD-Cy5. The E_FRET_ decreases in a monophasic manner down to the E_FRET_ trace predicted for no interaction between ddP/5′TCy3 DNA and RPA-OBA-Cy5 (**Figure 2B**, *Bottom*). This indicates that Rad51•ATP filaments irreversibly exchange with RPA on all ddP/5′TCy3 DNA, displacing all RPA into solution. The E_FRET_ decrease observed for RPA/Rad51 exchange is best fit to a one-step, irreversible kinetic model (i.e., single exponential decline) with a rate constant of *k*_obs dec_ = 8.59 ± 0.05 (× 10^−3^) s^−1^ (**Table 1**). Interestingly, RPA/Rad51 exchange observed in **Figure 2B** is ~2.2-fold slower compared to that observed for the reference condition where the P/T DNA•RPA complexes are devoid of PCNA (*k*_obs dec_ = 1.87 ± 0.01 (× 10^−2^) s^−1^, **Figure S3**, **Table 1**). Hence, the pre-occupancy of P/T DNA•RPA complexes by loaded PCNA slows but does not prohibit RPA/Rad51exchange.

**Table 1.** Kinetic rate constants observed in the present study for RPA/Rad51 exchange and RPA/Rad51 exchange-dependent PCNA unloading. N/A = Not Applicable.

| Figure |  | S3B | 2B | 2C | 3B | 3C | 5B | 5C |
| --- | --- | --- | --- | --- | --- | --- | --- | --- |
| Reaction Components | RPA | + **Cy5**-RPA | + **Cy5**-RPA | RPA | + **Cy5**-RPA | RPA | + **Cy5**-RPA | RPA |
|  | RFC, PCNA | - | + RFC, PCNA | + RFC, **Cy5**-PCNA | + RFC, PCNA | + RFC, **Cy5**-PCNA | + RFC, PCNA | + RFC, **Cy5**-PCNA |
| | Pol $\delta$ | - | - | - | + | + | + | + |
|  | dGTP | - | - | - | - | - | + | + |
|  | Rad51 | + | + | + | + | + | + | + |
| Rate Constant(s) | RPA/Rad51 Exchange | $1.87 \pm 0.01 \times (10^{-2}) \text{ s}^{-1}$ | $8.59 \pm 0.05 \times (10^{-3}) \text{ s}^{-1}$ | N/A | $1.23 \pm 0.01 \times (10^{-2}) \text{ s}^{-1}$ | N/A | $2.05 \pm 0.03 \times (10^{-2}) \text{ s}^{-1}$ | N/A |
| | RPA/Rad51 Exchange-dependent PCNA Unloading | N/A | N/A | $5.47 \pm 0.03 \times (10^{-3}) \text{ s}^{-1}$<br>$9.64 \pm 0.02 \times (10^{-4}) \text{ s}^{-1}$ | N/A | $4.19 \pm 0.3 \times (10^{-2}) \text{ s}^{-1}$<br>$9.16 \pm 0.2 \times (10^{-3}) \text{ s}^{-1}$<br>$1.16 \pm 0.05 \times (10^{-3}) \text{ s}^{-1}$ | N/A | Not Observed |

For experiments monitoring interactions of PCNA with P/T junctions (**Figure 2A**, *Right*), *I*_563_ and *I*_665_ values are stabile over time (**Figure 2D**, *Top*) and a significant, constant E_FRET_ is observed prior to the addition of Rad51•ATP (**Figure 2D**, *Bottom*) due to the loading and maintenance of Cy5-PCNA on all 5′ddPCy3/T DNA•RPA complexes^15^. Upon addition of Rad51•ATP, *I*_563_ increases simultaneously with decreases in *I*_665_ (**Figure 2D**, *Top*). These synchronized, anti-correlated changes in *I*_563_ and *I*_665_ are indicative of a direct FRET decrease due to the increase in distance between the Cy3 FRET donor on the 5′ddPCy3/T DNA and the Cy5 FRET acceptor on Cy5-PCNA. The E_FRET_ decreases in a biphasic manner down to the E_FRET_ trace predicted for no interaction between 5′ddPCy3/T DNA and Cy5-PCNA (**Figure 2D**, *Bottom*). This indicates that all PCNA encircling 5′ddPCy3/T DNA irreversibly unload. The observed E_FRET_ decrease is best fit to a two-step, irreversible kinetic model with rate constants of *k*_obs dec 1_ = 5.47 ± 0.03 (× 10^−3^) s^−1^ and *k*_obs dec 2_ = 9.64 ± 0.02 (× 10^−4^) s^−1^ where *k*_obs dec 2_ reports on PCNA unloading (**Table 1**). Direct comparisons of the data presented in **Figures 2B** and **2C** indicate that upon addition of Rad51•ATP both RPA/Rad51 exchange and PCNA unloading go to completion and are irreversible. Next, we sought to identify correlations between RPA/Rad51 exchange and the occupancy of P/T junctions by PCNA.

The E_FRET_ decreases observed in **Figures 2B** and **Figure 2C** were each normalized to their respective ranges and the results plotted as function of time (after addition of Rad51•ATP). As observed in **Figure S4**, PCNA begins unloading from P/T junctions immediately upon addition of Rad51•ATP and becomes significant (i.e., ≥ 10%) when RPA/Rad51 exchange ≥ 90% completed. ~40% of PCNA unloads as RPA/Rad51 exchange is occurring on P/T junctions; the remaining ~60 % unloads after RPA/Rad51 exchange has been completed on all P/T junctions (i.e., RPA has been replaced by Rad51•ATP filaments on all P/T junctions). Altogether, this reveals that unloading of PCNA from P/T DNA•RPA complexes requires RPA/Rad51 exchange but not vice versa. Rather, the occupancy of P/T DNA•RPA complexes by loaded PCNA slows but does not prohibit RPA/Rad51exchange (**Figures 2B**, **S3**, and **Table 1**).

### An abundant PCNA-binding protein complex does not stabilize loaded PCNA on P/T junctions during RPA/Rad51 exchange

Pol δ is maintained at stalled replication sites induced by UVR exposure and may re-engage the resident PCNA, forming holoenzymes that re-iteratively associate and dissociate from the P/T junctions. However, pol δ holoenzymes cannot support stable insertion of a dNTP opposite the offending lesion. Furthermore, hundreds of additional PCNA-binding proteins/protein complexes are maintained or enriched on PCNA encircling P/T junctions of stalled replication sites induced by UVR exposure^15,24,49–54^. To emulate RPA/Rad51 exchange in this scenario where PCNA-binding proteins/protein complexes are locally abundant, we repeated the experiments described above (in **Figure 2**) exactly as indicated except that PCNA encircling P/T DNA•RPA complexes is preincubated with a 2-fold excess of pol δ prior to the addition of Rad51•ATP (**Figure 3A**). Under these conditions, pol δ binding to loaded PCNA is stoichiometric and unloading of PCNA from P/T DNA•RPA complexes is not observed up to a 4-fold excess of pol δ^15^. Excess pol δ is utilized to ensure all PCNA encircling P/T DNA•RPA complexes are engaged in pol δ holoenzymes prior to the addition Rad51•ATP^15^. However, in the absence of dNTPs, pol δ holoenzymes have dramatically low affinity, if any, for native (i.e., undamaged) P/T junctions. Thus, pol δ holoenzymes do not stably engage P/T junctions in these experiments, emulating that observed at P/T junctions of stalled replication sites^4,15,50–53^.

**Figure 3.**
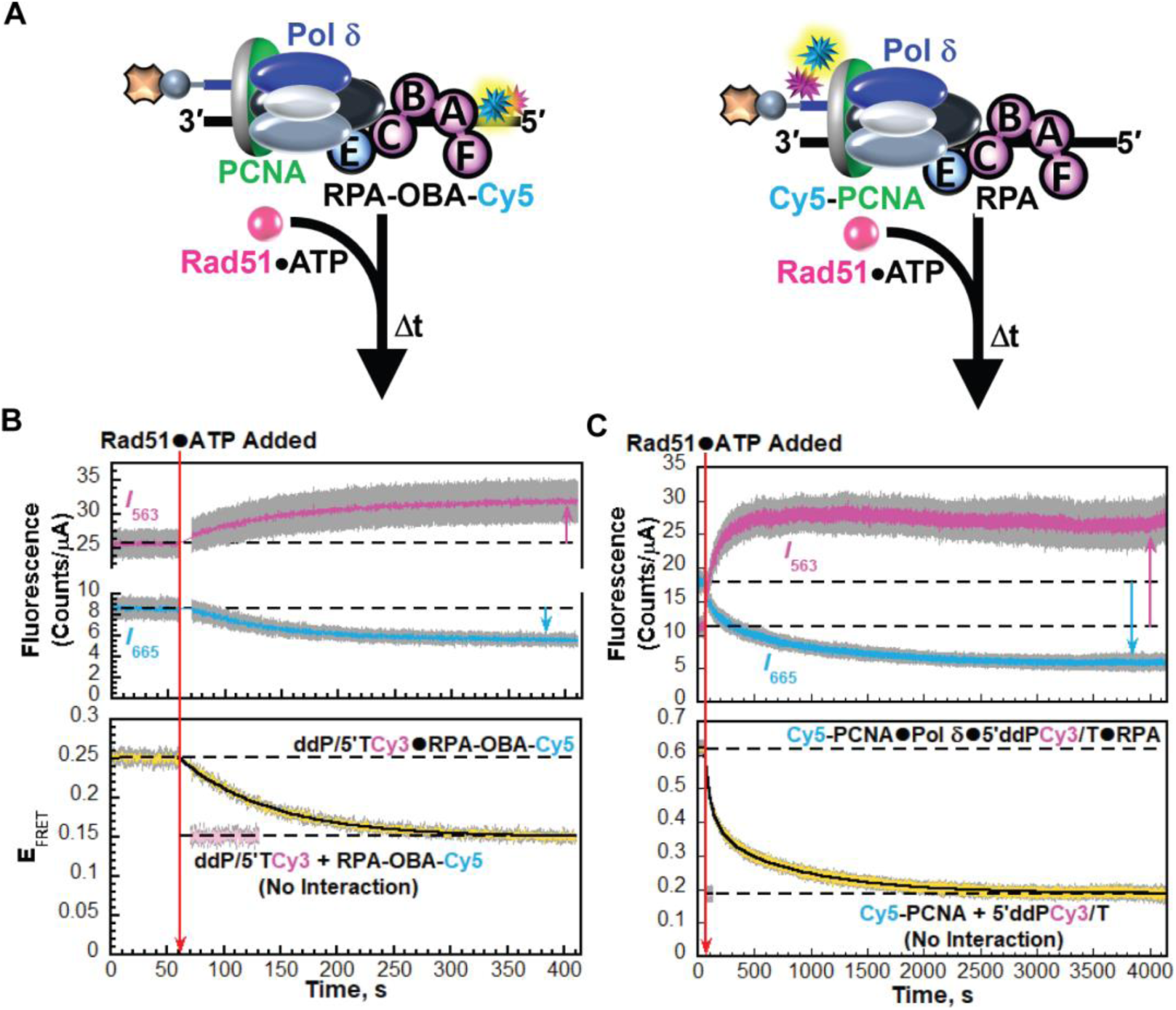
Pol δ does not stabilize loaded PCNA on P/T junctions during RPA/Rad51 exchange. (A) Schematic representations of the FRET pairs and experiments to monitor the interactions of RPA (Left) and PCNA (Right) with P/T junctions during RPA/Rad51 exchange in the presence of pol δ (B - C) Data. Each trace is the mean of at least three independent traces with the S.E.M. shown in grey. The time trajectories of I563 (magenta) and I665 (cyan) are displayed in the top panels and the corresponding EFRET (yellow) is displayed in the bottom panels. The times at which Rad51•ATP are added is indicated by red arrows. Changes in I563 and I665 are indicated in the top panels by magenta and cyan arrows, respectively. For observation, the I563 and I665, and the corresponding EFRET values observed prior to the addition of Rad51•ATP are each fit to dashed flat lines that are extrapolated to the axes limits. The predicted EFRET traces for no interaction between the respective Cyanine-labeled species are displayed in pink in each bottom panel, offset to the time after Rad51•ATP is added and fit to a dashed flat line that is extrapolated to the axis limits. Data for interactions of RPA and PCNA with P/T junctions during RPA/Rad51 exchange in the presence of pol δ are shown in panels B and C, respectively. The rate constant(s) for the EFRET decrease observed after the addition of Rad51•ATP in each bottom panel is reported in Table 1.

For experiments monitoring interactions of RPA with P/T junctions (**Figure 3A**, *Left*), the time-dependent traces for *I*_563_ and *I*_665_ (**Figure 3B**, *Top*) and E_FRET_ (**Figure 3B**, *Bottom*) are similar to those observed in **Figure 2B** where pol δ was absent. This indicates that Rad51•ATP filaments irreversibly exchange with RPA on all ddP/5′TCy3 DNA under these conditions, displacing all RPA into solution. The E_FRET_ decrease observed for RPA/Rad51 exchange is best fit to a one-step, irreversible kinetic model (i.e., single exponential decline) with a rate constant [*k*_obs dec_ = 1.23 ± 0.01 (× 10^−2^) s^−1^] that is ~1.5-fold slower compared to that observed for the reference condition (**Figure S3**, **Table 1**). Together with the observations described above, this indicates that occupancy of P/T DNA•RPA complexes by either PCNA alone (**Figure 2B**) or PCNA engaged in pol δ holoenzymes (**Figure 3B**) slows but does not prohibit RPA/Rad51exchange.

For experiments monitoring interactions of PCNA with P/T junctions (**Figure 3A**, *Right*), the behaviors of the time-dependent traces for *I*_563_ and *I*_665_ (**Figure 3C**, *Top*) and E_FRET_ (**Figure 3C**, *Bottom*) are similar to those observed in **Figure 2C** where pol δ was absent. This indicates that, despite being pre-engaged by in pol δ holoenzymes, all PCNA encircling ddP/5’TCy3 DNA irreversibly unload. Interestingly, the E_FRET_ decrease observed for PCNA unloading under these conditions (**Figure 3C**, *Bottom*) is best fit to a triple exponential decline with a fast rate constant (*k*_obs dec 1_ = 4.19 ± 0.3 (× 10^−2^) s^−1^) that accounts for 19.5% ± 0.8% of the decline and is significantly greater than the slower rate constants (*k*_obs dec 2_ = 9.16 ± 0.2 (× 10^−3^) s^−1^, *k*_obs dec 3_ = 1.16 ± 0.05 (× 10^−3^) s^−1^) (**Table 1**). The fast rate constant is not observed when pol δ is omitted (**Figure 2C**, *Bottom*). The slower rate constants (*k*_obs dec 2_ and *k*_obs dec 3_) observed in the presence of pol δ are very similar to the rate constants (*k*_obs dec 1,_ *k*_obs dec 2_) observed in the absence of pol δ (**Table 1**). Consequently, upon addition of Rad51•ATP, subsequent unloading of PCNA from P/T DNA is achieved relatively faster when pre-loaded PCNA is engaged in pol δ holoenzymes compared to when it is not (**Figure S5**). Thus, pol δ holoenzymes do not impart any stability to loaded PCNA during RPA/Rad51 exchange. This suggests that tight interactions of PCNA-binding proteins/protein complexes with loaded PCNA do not stabilize sliding clamps on P/T junctions during RPA/Rad51 exchange. Direct comparisons of the data presented in **Figures 3B** and **3C** indicate that upon addition of Rad51•ATP both RPA/Rad51 exchange and PCNA unloading go to completion and are irreversible under these conditions. Next, we compared the timescales of RPA/Rad51 exchange and PCNA unloading under the conditions depicted in **Figure 3A** to identify correlations between these two processes.

As observed in **Figure S6**, PCNA begins unloading from P/T junctions immediately upon addition of Rad51•ATP and becomes significant (i.e., ≥ 10%) when RPA/Rad51 exchange is only ~ 10 % completed. ~75% of PCNA unloads as RPA/Rad51 exchange is occurring on the P/T DNA and the remaining ~25 % unloads after RPA has been replaced by Rad51•ATP filaments on all P/T DNA. This contrasts that observed in the absence of pol δ where the majority (~60%) of PCNA unloads after RPA/Rad51 exchange has been completed on all P/T DNA (**Figure S4**). Regardless, this confirms that unloading of PCNA from P/T DNA•RPA complexes requires RPA/Rad51 exchange but not vice versa. Next, we investigate the mechanism(s) of RPA/Rad51 exchange-dependent PCNA unloading.

### RPA/Rad51 exchange on P/T DNA eliminates the resident PCNA by inhibiting RFC

In the experiments described in **Figures 2** and **3** above, P/T junctions are initially engaged by RPA and ATP-bound RFC (i.e., RFC•ATP) pre-loads PCNA onto all P/T junctions. After dissociation of the RFC•ADP complex into solution, loaded PCNA is left behind on the dsDNA region upstream of the P/T junctions^15,33^. The biotin-neutravidin complexes prohibit diffusion of loaded PCNA off the blunt duplex ends of the P/T DNA. RPA•ssDNA complexes prohibit diffusion of loaded PCNA off the ssDNA ends of the P/T DNA as well as RFC-catalyzed unloading of PCNA^4,37^. Thus, prior to the addition of Rad51•ATP, the only pathway for unloading of PCNA from DNA is through spontaneous opening of the PCNA ring. This non-catalytic process is dramatically slow and, in the event that it occurs, RPA permits RFC•ATP to instantly “re-load” free PCNA back onto P/T junctions such that unloading is not observed^2,4,15,24^. This is confirmed for the experimental system in the current study (**Figure S7**). Hence, the stable (i.e., equilibrium) E_FRET_ traces observed prior to the addition of Rad51•ATP in **Figures 2C** and **3C** represent RFC•ATP maintaining loaded Cy5-PCNA on all P/T DNA•RPA complexes.

In the present study, all rate constants observed for RPA/Rad51 exchange-dependent PCNA unloading are ≥ ~13-fold slower than the rate constant for RFC-catalyzed unloading of PCNA (*k*_obs_ = 0.54 ± 0.09 s^−1^)^2^. This indicates that RPA/Rad51 exchange on P/T junctions does not promote RFC-catalyzed unloading of the resident PCNA. Furthermore, ATP hydrolysis by Rad51 is restricted in the present study, Rad51 lacks any known PCNA-binding motif(s), and, to the best of our knowledge, direct interactions between Rad51 and PCNA have yet to be reported. This indicates that Rad51 does not catalyze PCNA unloading. Altogether, this suggests that RPA/Rad51 exchange-dependent PCNA unloading is non-catalytic. Two general populations are observed for this process. The “slow” population is ≤ 2.0-fold slower than spontaneous opening of the PCNA ring (*k*_dec,obs_ = 1.93 ± 0.01 (× 10^−3^) s^−1^, **Figure S7**) and accounts for 100% and 44.3 ± 1.1 % of the unloading observed in **Figure 2C** (*k*_obs dec 2_) and **Figure 3C** (*k*_obs dec 3_), respectively. The “fast” population is ≥ ~ 4.74-fold faster than spontaneous opening of the PCNA ring, accounts for 55.7 ± 1.1 % of the unloading observed in **Figure 3C** (*k*_obs dec 1_, *k*_obs dec 2_), and only appears when Rad51•ATP is added to loaded PCNA pre-assembled in pol δ holoenzymes. Altogether, this suggests that RPA/Rad51 exchange-dependent PCNA unloading is non-catalytic and occurs through spontaneous opening of the PCNA ring or a mechanism where Rad51•ATP and pol δ in conjunction somehow destabilize the PCNA ring.

RPA/Rad51-dependent PCNA unloading observed in **Figures 2B** and **2**C generates “free” PCNA in solution, goes to completion, and is irreversible, despite the continuous presence of RFC•ATP. This is only possible if assembly of stable Rad51•ATP filaments on P/T junctions via RPA/Rad51 exchange also prohibits RFC•ATP from loading free PCNA onto P/T junctions that are devoid of the sliding clamp. To confirm this, we monitor and directly compare RFC-catalyzed loading of “free” PCNA on P/T junctions engaged by either RPA or Rad51•ATP filaments (**Figure 4**)^15^. 5′ddPCy3/T DNA substrates are first pre-bound with NeutrAvidin and either RPA or Rad51•ATP filaments. Next, Cy5-PCNA is added and the resultant solution is equilibrated. Finally, RFC•ATP is added. For experiments on P/T junctions engaged by RPA (**Figure 4A**, *Left*), *I*_563_ and *I*_665_ values are stabile over time (**Figure 4B**, *Top*) and a low, constant E_FRET_ is observed prior to the addition of RFC•ATP (**Figure 4B**, *Bottom*) due to the presence of both the Cy3 donor and the Cy5 acceptor. E_FRET_ values observed during this period are a true experimental baseline representing the absence of any interactions between Cy5-PCNA and the 5′ddPCy3/T DNA•RPA complexes^15^. Upon addition of RFC•ATP, *I*_665_ increases concomitantly with a decrease in *I*_563_ after which both fluorescence emission intensities stabilize and persist (**Figure 4B**, *Top*). These synchronized, anti-correlated changes in *I*_563_ and *I*_665_ are direct indicators of the appearance and increase in FRET (**Figure 4B**, *Bottom*) due to the decrease in distance between the Cy3 FRET donor on the 5′ddPCy3/T DNA substrate and the Cy5 FRET acceptor on Cy5-PCNA. The observed E_FRET_ increase (**Figure 4B**, *Bottom*) is biphasic with *k*_obs inc 1_ = 4.46 ± 0.2 (× 10^−2^) s^−1^ and *k*_obs inc 2_ = 1.70 ± 0.12 (× 10^−2^) s^−1^. These rate constants are in excellent agreement with values obtained in identical experiments from a recent study^15^. At the plateau of the E_FRET_ increase, Cy5-PCNA is loaded and maintained on all 5′ddPCy3/T•RPA complexes^15^.

**Figure 4.**
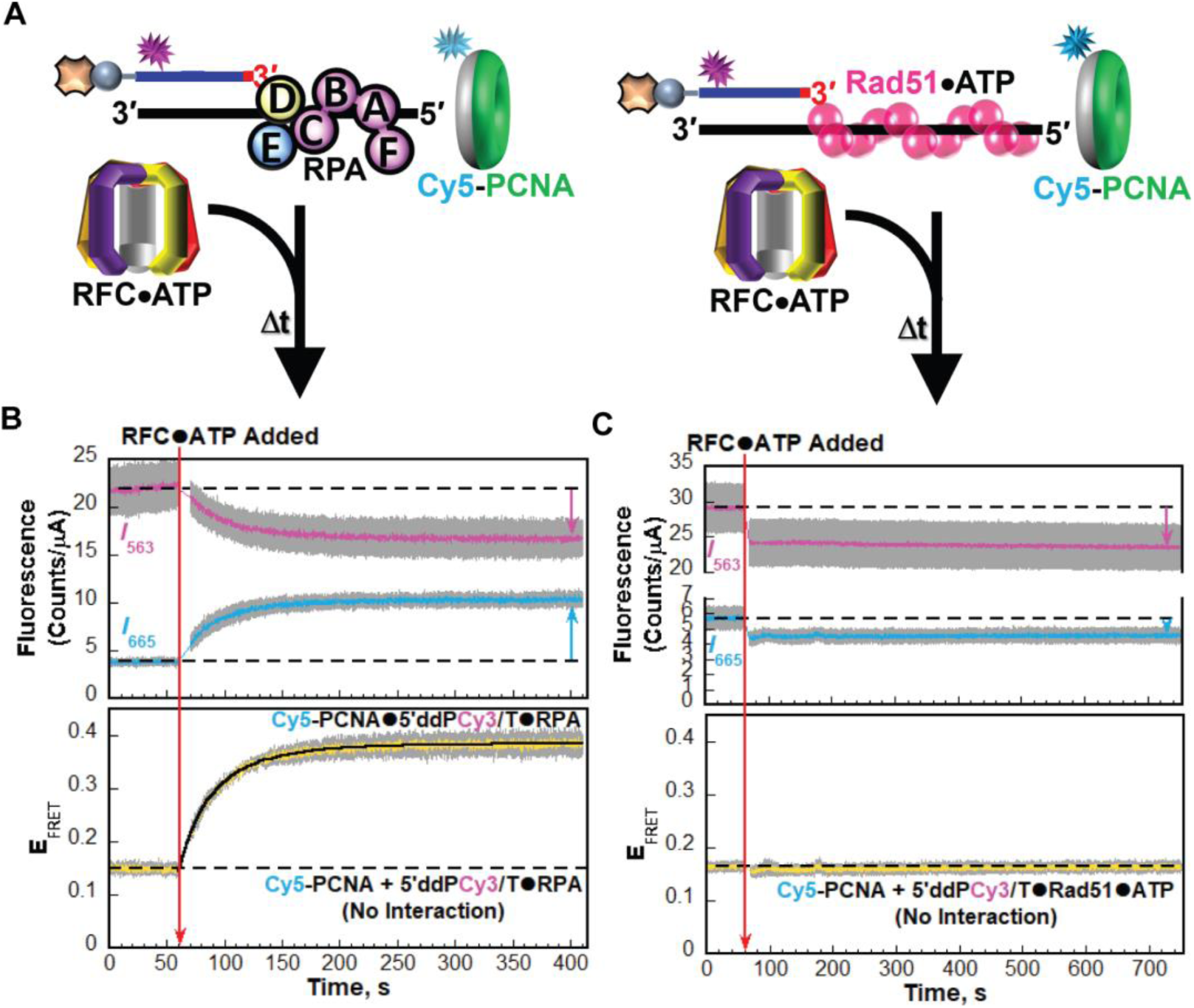
Rad51•ATP filaments prohibit RFC-catalyzed loading of PCNA onto P/T junctions. (A) Schematic representations of the FRET pair and experiments to monitor the RFC-catalyzed loading of PCNA onto P/T junctions engaged by RPA (*Left*) or Rad51•ATP filaments (*Right*) (**B - C**) Data. Each trace is the mean of at least three independent traces with the S.E.M. shown in grey. The time trajectories of *I*563 (magenta) and *I*665 (cyan) are displayed in the top panels and the corresponding EFRET (mustard) is displayed in the bottom panels. The time at which RFC•ATP is added is indicated by a red arrow. Changes in *I*563 and *I*665 are indicated in the top panel by magenta and cyan arrows, respectively. For observation, the *I*563 and *I*665, and the corresponding EFRET values observed prior to the addition RFC•ATP are each fit to dashed flat lines that are extrapolated to the axis limits. (**B**) Data for RFC-catalyzed loading of PCNA onto P/T junctions engaged by RPA. (**C**) Data for RFC-catalyzed loading of PCNA onto P/T junctions engaged by Rad51•ATP filaments.

For experiments on P/T junctions engaged by Rad51•ATP filaments (**Figure 4A**, *Right*), *I*_563_ and *I*_665_ values are stabile over time (**Figure 4C**, *Top*) and a low, constant E_FRET_ is observed prior to the addition of RFC•ATP (**Figure 4C**, *Bottom*) due to the presence of both the Cy3 donor and the Cy5 acceptor. Again, E_FRET_ values observed during this period are a true experimental baseline representing the absence of any interactions between Cy5-PCNA and the 5′ddPCy3/T DNA•Rad51•ATP complexes. Upon addition of RFC•ATP, both *I*_665_ and *I*_563_ slightly but reproducibly decrease within the dead time after which both *I* values remain flat (**Figure 4B**, *Top*). These synchronized, correlated changes in *I*_563_ and *I*_665_ indicate that any observed changes in FRET are indirect^15,55^. Nevertheless, a change in *E*_FRET_ is not observed upon addition of RFC•ATP due to *I*_563_ and *I*_665_ each decreasing by the same relative amount (*I*_665_ by 10.6 ± 3.1 % and *I*_563_ by 10.3 ± 2.0 %). This confirms that assembly of stable Rad51•ATP filaments on P/T junctions via RPA/Rad51 exchange prohibit RFC•ATP from loading “free” PCNA on P/T junctions that are devoid of the sliding clamp.

### Rad51•ATP does not require access to 3′ primer termini of P/T junctions and the ssDNA immediately downstream for RPA/Rad51 exchange

Formation of Rad51•ATP filaments occurs in a stepwise in a manner. First, 1 - 3 Rad51 monomers, each engaged with ATP, serve as a nucleus to initiate filament assembly (i.e., nucleate) on a template ssDNA. Next, the filament nuclei extend (i.e., grow/elongate) in either direction by the addition of Rad51 monomers, each engaged with ATP^46–48^. Previous single molecule studies on a “gapped” DNA substrate comprised of dsDNA regions (i.e., handles) connected by an RPA-coated ssDNA gap revealed that nucleation of Rad51 is intrinsically enriched at/near the 3′ dsDNA/ssDNA junction compared to the 5′ dsDNA/ssDNA junction^47^. A 3′ dsDNA/ssDNA junction contains a recessed 3′ primer terminus (from where DNA synthesis can initiate from) and a 5′ template ssDNA overhang. These junctions are amenable to those utilized in the present study (i.e., P/T junctions). Thus, we next sought to address whether Rad51•ATP requires access to the recessed 3′ primer termini of P/T junctions for nucleation and, hence, RPA/Rad51 exchange. To do so, we repeated the experiments described above in **Figure 3** above except that loaded PCNA is preincubated with a 2-fold excess of pol δ and dGTP prior to the addition of Rad51•ATP (**Figure 5A**). Under these conditions, pol δ engages the “front face” of loaded PCNA, forming a holoenzyme, engages the P/T junction, and aligns an incoming dGTP at the 3′ terminus of the primer strand in a complimentary bp with the template nucleotide (a Cytosine) immediately 5′ of the P/T junction. However, both extension of the primer (via pol δ DNA polymerase activity) and degradation of the primer (via pol δ exonuclease activity) are prohibited due to the 3′ dideoxy-terminated primer strand and the use of exonuclease-deficient pol δ. Hence, the pol δ holoenzymes are trapped in initiation states for DNA synthesis. In this state (depicted in **Figure 5A**), RPA has vacated the 3′ primer termini of P/T junctions and ~ 6 nts of the template ssDNA immediately downstream and these sites are sequestered within the DNA polymerase active site of pol δ^4,15,52,56^.

**Figure 5.**
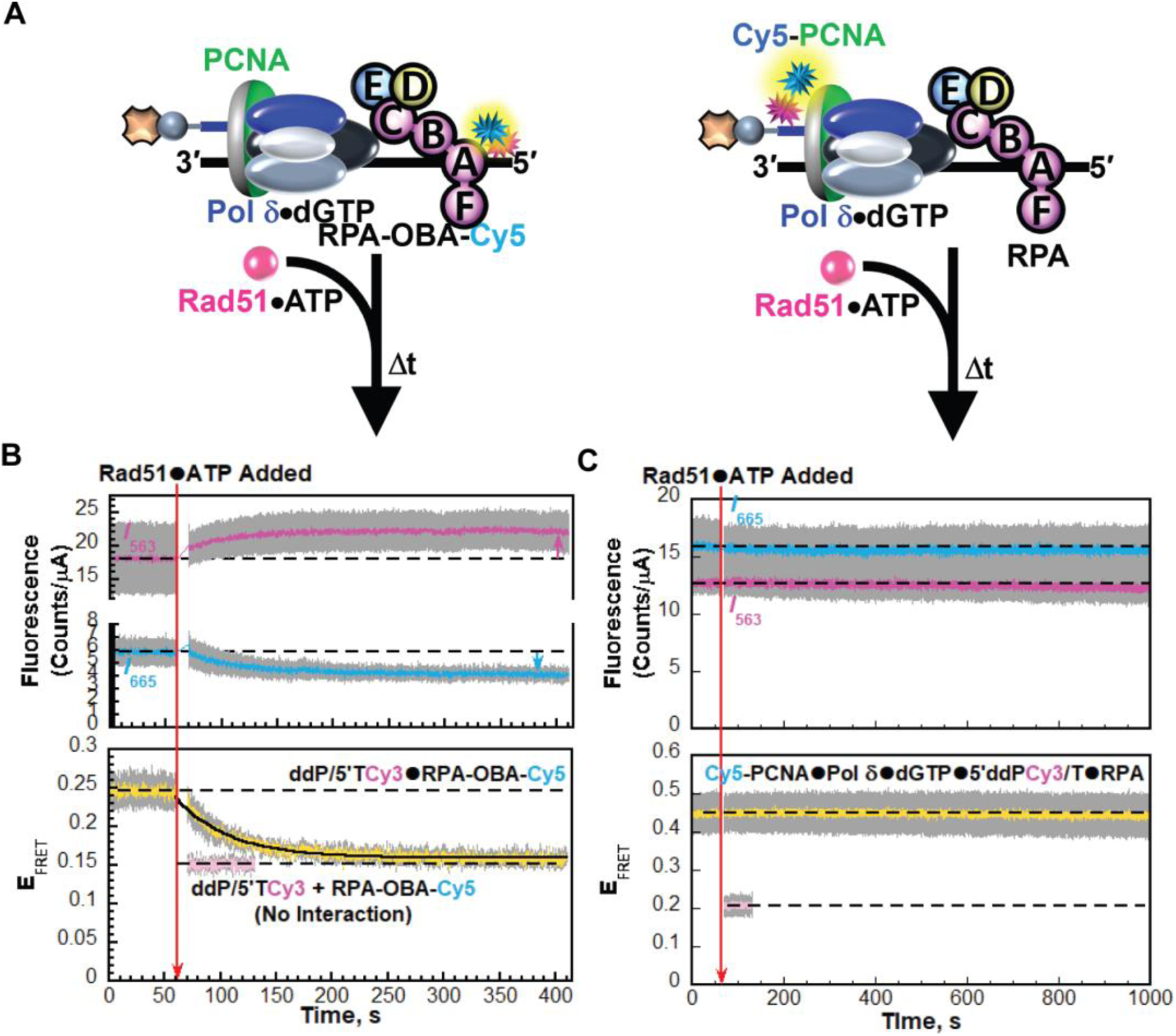
Rad51•ATP does not require access to 3’ primer termini of P/T junctions and the ssDNA immediately downstream for RPA/Rad51 exchange. (A) Schematic representations of the FRET pairs and experiments to monitor the interactions of RPA (*Left*) and PCNA (*Right*) with P/T junctions during RPA/Rad51 exchange in the presence of pol δ and dGTP (**B - C**) Data. Each trace is the mean of at least three independent traces with the S.E.M. shown in grey. The time trajectories of *I*563 (magenta) and *I*665 (cyan) are displayed in the top panels and the corresponding EFRET (mustard) are displayed in the bottom panels. The times at which Rad51•ATP is added are indicated by red arrows. Changes in *I*563 and *I*665, if they occur, are indicated in the top panels by magenta and cyan arrows, respectively. For observation, the *I*563 and *I*665, and the corresponding EFRET values observed prior to the addition Rad51•ATP are each fit to dashed flat lines that are extrapolated to the axis limits. The predicted EFRET traces for no interaction between the respective Cyanine-labeled species are displayed in pink in each bottom panel, offset to the time after Rad51•ATP is added and fit to a dashed flat line that is extrapolated to the axis limits. Data for interactions of RPA and PCNA with P/T junctions during RPA/Rad51 exchange in the presence of pol δ and dGTP are shown in panels **B** and **C,** respectively. The rate constant for the EFRET decrease observed after the addition of Rad51•ATP in panel **B** is reported in **Table 1**.

For experiments monitoring interactions of RPA with P/T junctions (**Figure 5A**, *Left*), the time-dependent traces for *I*_563_ and *I*_665_ (**Figure 5B**, *Top*) and E_FRET_ (**Figure 5B**, *Bottom*) are similar to those observed in **Figures 2B** and **3B** where either dGTP was absent (**Figure 3B**) or both pol δ and dGTP were absent (**Figure 2B**). This indicates that Rad51•ATP filaments irreversibly exchange with RPA on all ddP/5′TCy3 DNA under these conditions, displacing all RPA into solution. Again, the E_FRET_ decrease observed during RPA/Rad51 exchange is best fit to a one-step, irreversible kinetic model with a rate constant of *k*_obs dec_ = 2.05 ± 0.03 (× 10^−3^) s^−1^, which is faster than all rate constants observed for RPA/Rad51 exchange in the current study (**Table 1**). For experiments monitoring interactions of PCNA with P/T junctions (**Figure 5A**, *Right*), *I*_563_ and *I*_665_ values are stabile over time (**Figure 5C**, *Top*) and a significant, constant E_FRET_ is observed prior to the addition of Rad51•ATP (**Figure CD**, *Bottom*) due to the initiation states of pol δ holoenzymes^15^. Upon addition of Rad51•ATP, the *I*_563_ and *I*_665_ traces are maintained (**Figure 5C**, *Top*) and a change in E_FRET_ is not observed for over 15 mins (**Figure 5C**, *Bottom*). This confirms that the initiation state of pol δ holoenzymes is achieved in these experiments prior to the addition of Rad51•ATP and reveals that this state stabilizes loaded PCNA on P/T junctions during RPA/Rad51 exchange (for ≥ 1000 s). Direct comparisons of the data presented in **Figures 5B** and **5C** indicate that Rad51•ATP nucleates and completely exchanges with RPA (**Figure 5B**) while the 3′ primer termini of the P/T junctions and the template ssDNA immediately downstream are sequestered by pol δ holoenzymes in the initiation state (**Figure 5C**). Thus, Rad51•ATP does not require access to the recessed 3′ primer termini of P/T junctions for nucleation and, hence, RPA/Rad51 exchange.

## Discussion

Upon the stalling of canonical DNA replication at DNA lesions, such as UVR-induced photoproducts, the resident PCNA are left behind on the dsDNA regions upstream of the afflicted P/T junctions, which are directly engaged by RPA (**Figure 6**, *Top Left*). RPA also coats the unreplicated ssDNA templates downstream^19,20,57^. In humans, these stalling events are circumvented by one of at least three interconnected pathways; TLS, TS, HDR. PCNA encircling P/T junctions of stalled replication sites are central to the activation of both TLS and TS (**Figure 1B**). As described above, “protein roadblocks” restrict diffusion of loaded PCNA on DNA, collectively localizing the sliding clamps at/near P/T junctions of stalled replication sites^2,4,15,23,24,33–37,58,59^. Nevertheless, PCNA can unload from these sites and this process can be catalyzed by enzyme complexes, such as ATAD5, or occur via spontaneous opening of the PCNA ring (**Figure S7**)^2,60–65^. For loading PCNA onto P/T junctions, assembled RFC•ATP•PCNA complexes, i.e., loading complexes, require direct interactions with the 3′ termini of primer strands and the template ssDNA immediately downstream^52,66^. RPA directly engages these sites but permits access to loading complexes through microscopic dissociation events within the RPA complex. Consequently, RFC loads and maintains PCNA on P/T junctions that are directly engaged by RPA (**Figure S7**)^4,15^. This suggests that at stalled replication sites where the P/T junctions are directly engaged by RPA, RFC maintains loaded PCNA at these sites, promoting TLS and/or TS (**Figure 6**, *Left*).

**Figure 6.**
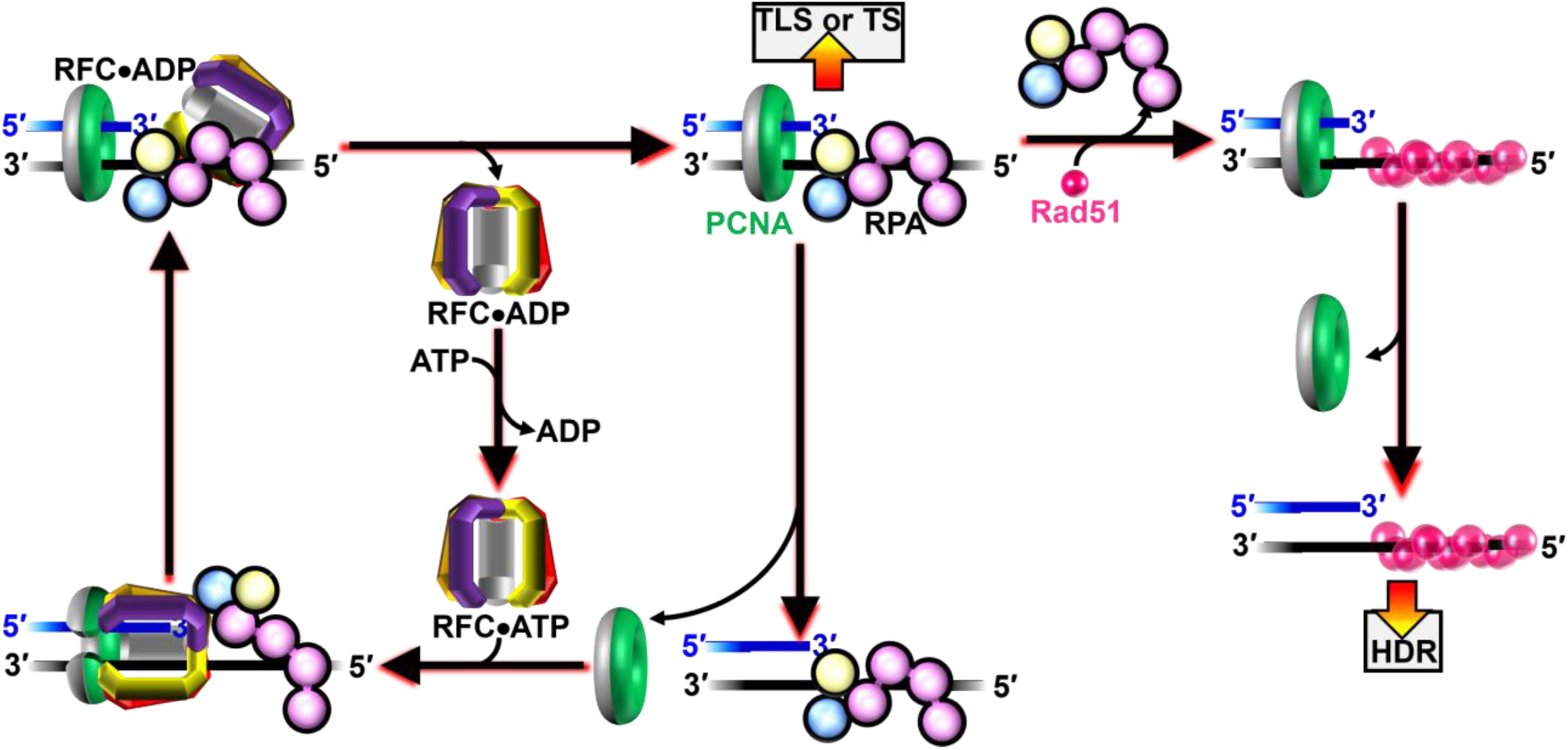
Model for interplay between RPA/Rad51 exchange and PCNA occupancy at P/T junctions of stalled replication sites. (*Left*) RFC maintains PCNA on P/T junctions engaged by RPA, promoting TLS and/or TS. (*Right*) Assembly of stable Rad51·ATP filaments on P/T junctions of stalled replication sites via RPA/Rad51 exchange ultimately leads to the complete elimination of the resident PCNA, prohibiting TLS and TS and promoting HDR.

Exchange of RPA for Rad51•ATP along the unreplicated ssDNA template sequences of stalled replication sites is central to the activation of HDR (**Figure 1B**)^21,22,32,67,68^. The results from the present study reveal that PCNA encircling P/T junctions engaged by RPA unload slowly via non-catalytic mechanisms during and after (the complete) exchange of RPA for Rad51•ATP on P/T junctions (**Figures 2-3**, **S4**, **S6**, **S7**). Furthermore, assembly of stable Rad51•ATP filaments on P/T junctions via complete RPA/Rad51 exchange prohibit RFC•ATP from loading “free” PCNA back onto P/T junctions that are devoid of the sliding clamp (**Figure 4**). Together, this leads to complete elimination of loaded PCNA from P/T junctions. This phenomenon is observed both in the absence and presence of saturating pol δ, an abundant, high-affinity PCNA-binding protein complex that is maintained at stalled replication sites induced by UVR exposure (**Figures 2-3**, **S4**-**S6**)^49,54^. Multiple studies from many independent groups revealed that stable Rad51•ATP filaments directly and statically engage the 3′ primer termini of P/T junctions and the template ssDNA immediately downstream and these interactions effectively shelter (i.e., “protect”) the 3′ primer termini from processing by DNA metabolic enzymes, such as 3′5′ exonucleases^69^. Thus, stable Rad51•ATP filaments on P/T junctions prohibit PCNA loading (as observed in **Figure 4B**) by restricting access of loading complexes to 3′ primer termini of P/T junctions and template ssDNA immediately downstream. Altogether, this suggests that assembly of stable Rad51•ATP filaments at P/T junctions of stalled replication sites via RPA/Rad51 exchange ultimately leads to the complete elimination of the resident PCNA, prohibiting TLS and TS and promoting HDR (**Figure 6**, *Right*).

According to the model proposed in **Figure 6**, upon stalling of canonical replication at UVR-induced lesions, the occupancy of the afflicted P/T junctions by PCNA, RPA, and Rad51•ATP are anti-correlated temporally, leading to a transition from TLS/TS to HDR. This agrees with results from recent iPOND studies of human cells treated with UVR. Here, as stalling prolonged at UVR-induced lesions, the levels of PCNA, RPA, and pol η at P/T junctions of stalled replications sites decreased over time as Rad51 levels increased at these sites^19^. Pol η is the TLS pol primarily responsible for the bypass of UVR-induced lesions via TLS. These and other observations led the authors to conclude a chronology of DDT pathways at UVR-stalled replication sites where TLS is the favored, early DDT pathway elicited. The results from the present study support this model.

The unreplicated ssDNA template sequences of UVR-stalled replication sites can reach 200 – 450 nts in length in human cells, which equates to RPA•ssDNA filaments comprised of ~6 - 15 RPA heterotrimers each in the 30 nucleotide binding mode^5–7,19,20,57,70^. Human Rad51 has an intrinsic preference for nucleating at/near P/T junctions in the presence of RPA but does not require access to these sites for nucleation and subsequent RPA/Rad51 exchange (**Figure 5**)^47^ ^47^. Nevertheless, Rad51•ATP filaments must ultimately engage the P/T junctions via RPA/Rad51 exchange to eliminate the resident PCNA (**Figure 6**, *Right*). *In vivo*, other cellular factors likely influence the temporal localization of RPA, PCNA, and Rad51 at the P/T junctions of stalled replication sites. For example, BRCA2, a molecular chaperone for Rad51 that accelerates filament nucleation, enhances Rad51•ATP deposition at dsDNA/ssDNA junctions in the presence of RPA^47,71^. Such activity at UVR-stalled replication sites may promote elimination of PCNA from the afflicted P/T junctions via inhibition of RFC (**Figure 4**). Furthermore, a recent study of human cells treated with hydroxyurea (HU) discovered that the PCNA unloader ATAD5 promotes interactions of Rad51 with P/T junctions of HU-stalled replication sites by catalytically unloading PCNA from these sites and subsequently recruiting Rad51 via direct protein•protein interactions. Hydroxyurea stalls canonical replication by depleting cellular dNTP pools^72^. At UVR-stalled replication sites, these ATAD5 activities may collectively eliminate PCNA from the afflicted P/T junctions by catalytically unloading PCNA from these sites and promoting Rad51-dependent inhibition of RFC (**Figure 4**). Also, recent studies have identified RadX, a negative regulator of Rad51 that promotes filament disassembly by dual mechanisms. RadX binds to and oligomerizes on the ends of growing Rad51•ATP filaments, blocking filament elongation. Also, RadX directly and selectively interacts with Rad51•ATP filaments, stimulating ATP hydrolysis and dismantling filaments^73,74^. At UVR-stalled replication sites, these RadX activities may stall and/or reverse RPA/Rad51 exchange at the afflicted P/T junctions, promoting occupancy of these sites by loaded PCNA (**Figure 4**). These and other hypotheses are currently under investigation.

## SUPPLEMENTARY INFORMATION

Supplemental Information includes Supplementary Results, Supplementary Methods, and Supplementary Figures S1 – S7.

## ACKNOWLEDGEMENTS

We would like to thank all members of the Hedglin and Antony labs for their efforts in reviewing/proofreading the current manuscript. This work was supported by funding from the National Institutes of Health to S.K. (F99CA274696), E.A. (R01 GM130756, R01 GM133967, R35 GM149320, and S10 OD030343) and M.H. (R35 GM147238-02),

## CONFLICTS OF INTEREST

The authors declare that they have no conflicts of interest with the contents of this article.^1^

## AUTHOR CONTRIBUTIONS

J.L.N., G.Y., K.G.P., R.L.D., S.P., and V.K. expressed, purified, and characterized all proteins. J.L.N., L.O.R., and G.Y. performed the experiments. M.H., J.L.N., and G.Y. designed the experiments. J.L.N., G.Y., M.H. and E.A. analyzed the data and wrote the paper.

## Supplemental Information

### Supplementary Results

*Rad51 has a strong, intrinsic preference for forming filaments on ssDNA at physiological ionic strength.* Human Rad51 binds both ssDNA and dsDNA to form right-handed helical nucleoprotein filaments where each Rad51 monomer binds either 3 bp (for dsDNA) or 3 nt (for ssDNA) ^1–3^. However, in the absence of ATP hydrolysis, Rad51•ATP has an exceptionally strong, intrinsic preference for forming filaments on ssDNA at physiological ionic strength (200 mM I) such that dsDNA binding is dramatically reduced, if observed at all. Binding of Rad51•ATP to dsDNA is only significant at low ionic strength^4–6^. To confirm that the specificity of Rad51•ATP for ssDNA at physiological ionic strength is achieved in the present studies we carried out Rad51 titrations on a P/T DNA substrate (FRET P/T, **Figure S1**) in which the ssDNA sequence is labeled with a Cy3 FRET donor and a Cy5 FRET acceptor. The former is located 3 nt downstream of the P/T junction and the latter is located 3 nt from the 5′ terminus of the ssDNA. The FRET P/T DNA substrate is comprised of a 15 bp duplex region that can accommodate 5 Rad51•ATP monomers and a 30 nt ssDNA region that can accommodate 10 Rad51•ATP monomers. In isolation, the ssDNA template of the FRET P/T DNA substrate forms a collapsed, flexible structure, bringing the two cyanine fluorophores close together and yielding a high E_FRET_ (**Figure S2A**). Stable interactions of Rad51•ATP with the ssDNA extends the engaged sequence ~1.6-fold relative to the B-form, thereby increasing the Cy3–Cy5 distance and reducing E_FRET7_. Thus, only saturation of the ssDNA sequence at increased concentrations of Rad51•ATP minimizes E_FRET_. The titration data is plotted as a function of the ratio of the amount Rad51 to the Total Amount Rad51-binding sites (i.e., dsDNA-binding sites + ssDNA-binding sites). Under the conditions of the assay, E_FRET_ decreases until the ssDNA sequence is saturated with Rad51•ATP, after which E_FRET_ remains constant at a minimal value. The fraction of Rad51 binding sites that are ssDNA is 0.66 (Rad51 ssDNA-binding sites/Total Rad51 binding sites = 10/15 = 0.667). Thus, if Rad51•ATP binds exclusively to ssDNA, the ssDNA sequence will be saturated with Rad51•ATP and the E_FRET_ will minimize at an equivalence point of Rad51:Total Rad51•ATP Binding Sites ~ 0.667. On the contrary, if Rad51•ATP engages both dsDNA- and ssDNA-binding sites indiscriminately, the ssDNA sequence will be saturated with Rad51•ATP and the E_FRET_ will minimize only when all Rad51 binding sites are saturated, i.e., at an equivalence point of Rad51:Total Rad51•ATP Binding Sites ~ 1.0. At physiological ionic strength (200 mM I), E_FRET_ decreases linearly and is minimized at an equivalence point of Rad51:Total Rad51-binding sites = 0.673 ± 0.014 (**Figure S2B**). This indicates that binding of Rad51•ATP to the ssDNA sequence of the FRET P/T DNA substrate is stoichiometric at 200 mM I, despite the presence of 15 bp of dsDNA. However, at low ionic strength (88 mM I), E_FRET_ decreases linearly and is minimized at an equivalence point of essentially Rad51:ssDNA Rad51-binding sites = 1.0 (0.981 ± 0.096, **Figure S2C**). This indicates that Rad51•ATP binds indiscriminately to dsDNA and ssDNA at low ionic strength. Altogether, this agrees with the results from previous studies and confirms that Rad51•ATP has an exceptionally strong, intrinsic preference for forming filaments on ssDNA over dsDNA under the experimental conditions of the current study (i.e., physiological ionic strength).

*RFC•ATP complexes maintain loaded PCNA on all P/T DNA engaged by RPA.* RPA engaged with P/T junctions permit RFC•ATP complexes to instantly “re-load” non-catalytically unloaded PCNA back onto P/T DNA such that the loss of PCNA is not observed over time^8–11^. Thus, RFC•ATP complexes continuously maintain loaded PCNA on all P/T junctions engaged by RPA. To confirm that this is established prior to the addition of Rad51•ATP complexes in the experiments described in **Figures 2C** and **3C** in the main text, we continuously monitor interactions of PCNA with P/T junctions engaged by RPA as reaction conditions progressively evolve (**Figure S7A**). First, 5′ddPCy3/T DNA is pre-bound with Neutravidin and native RPA. Next, Cy5-PCNA is added followed by RFC•ATP complexes and the fluorescence emission intensities are monitored over time until PCNA loading reaches equilibrium where PCNA is loaded and stabilized on all P/T junctions^8,9^. Under the conditions of the assay, RFC•ATP engages the “front face” of a free PCNA (in solution), opens the clamp, and the resultant complex, referred to as the loading complex, engages a P/T junction such that the “front face” of PCNA is oriented towards the 3′ terminus of the primer ^12^. Upon engaging a P/T junction, the loading complex adopts an activated conformation in which ATP hydrolysis by RFC is optimized. ATP hydrolysis by RFC simultaneously closes PCNA around the P/T junction and the closed (i.e., loaded) PCNA is subsequently released onto the dsDNA region of the P/T junction. The resultant RFC•ADP complex then releases into solution via dissociation from the resident RPA and subsequently exchanges ADP for ATP, re-forming RFC•ATP ^8,11–13^. In this scenario, PCNA loading is biphasic. First, all steps up to and including release of loaded PCNA onto the P/T junction are rate-limited by a kinetic step (*k*_obs inc 1_) along the pathway that occurs prior to and much slower than binding of loading complexes to P/T junctions. Second, loaded PCNA repositions relatively slowly (*k*_obs inc 2_) on the dsDNT region of the P/T junctions concurrent with release of RFC•ADP complexes into solution^8,9^. Thus, *k*_obs inc 1_ directly reports on the kinetics of loading “free” PCNA onto P/T junctions engaged by RPA. Once PCNA loading has reached equilibrium where PCNA is loaded and stabilized on all P/T junctions, unlabeled, native PCNA is added in excess and the fluorescence emission intensities are monitored over time. Under these conditions, the only pathway for unloading of Cy5-PCNA from P/T junctions is through spontaneous opening of the Cy5-PCNA ring. Once Cy5-PCNA is unloaded from the Cy3-labeled P/T DNA, re-loading of “free” Cy5-PCNA is prohibited due to the 70-fold excess of unlabeled, native PCNA. Consequently, the observed FRET decreases. Here, the disappearance of FRET is rate-limited by and directly reports on spontaneous opening of the PCNA ring.

Upon addition of RFC•ATP, *I*_665_ increases concomitantly with decreases in *I*_563_ after which both fluorescence emission intensities stabilize and persist (Figure S7B, *Top*). These synchronized, anti-correlated changes in *I*_563_ and *I*_665_ are indicative of the appearance and increase in FRET (Figure S7B, *Bottom*). As expected, the increase in E_FRET_ observed upon addition of RFC•ATP (Figure S7B, *Bottom*) is biphasic with *k*_obs inc 1_ = 4.46 + 0.2 (× 10^−2^) s^−1^ and *k*_obs inc 2_ = 1.70 + 0.12 (× 10^−2^) s^−1^. These rate constants are in excellent agreement with values obtained in identical experiments from a recent study^9^. The increase in E_FRET_ observed upon addition of RFC•ATP plateaus at values significantly above the E_FRET_ traces observed for no interaction between Cy5-PCNA and the 5′ddPCy3/T•RPA complexes. At this point, Cy5-PCNA is loaded and stabilized on all 5′ddPCy3/T•RPA complexes and RFC•ADP complexes have released into solution and exchanged ADP for ATP. Upon addition of excess, unlabeled PCNA, *I*_665_ decreases concomitantly with an increase in *I*_563,_ after which both fluorescence emission intensities stabilize and persist over time (Figure S7B, *Top*). These synchronized, anti-correlated changes in *I*_563_ and *I*_665_ are indicative of a decrease in FRET (Figure S7B, *Bottom*). The E_FRET_ decrease observed upon addition of excess, unlabeled PCNA bottoms out at E_FRET_ values observed for no interaction between Cy5-PCNA and the 5′ddPCy3/T•RPA complexes (i.e., all Cy5-PCNA remaining “free” in solution) indicating that all loaded Cy5-PCNA has non-catalytically unloaded from all 5′ddPCy3/T•RPA complexes (Figure S7B, *Bottom*).

Furthermore, the observed decrease E_FRET_ is comprised of a single phase (i.e., monophasic) with an observed rate constant [*k*_dec,obs_ = 1.93 + 0.01 (× 10^−3^) s^−1^] that is in good agreement with the rate constant for spontaneous opening of the PCNA ring [*k*_open_ = 1.25 + 0.32 (× 10^−3^) s^−1^]^14^. This confirms that dissociation of PCNA from P/T junctions engaged by RPA is governed entirely by spontaneous opening of the PCNA ring. Furthermore, the observed rate constant for RFC-catalyzed loading of “free” PCNA on P/T junctions engaged by RPA (*k*_obs inc 1_ = 4.46 + 0.2 (× 10^−2^) s^−1^) is ~23-fold faster than the observed rate constant for non-catalytic unloading of PCNA from P/T DNA (*k*_dec,obs_ = 1.93 + 0.01 (× 10^−3^) s^−1^). Hence, upon non-catalytic unloading of Cy5-PCNA from P/T junctions engaged by RPA, RFC•ATP complexes instantly reload Cy5-PCNA back onto the P/T junctions such that the loss of loaded Cy5-PCNA from 5′ddPCy3/T•RPA complexes is not observed^8,11^. In other words, for the experiments described in Figures 2 and 3 in the main text, RFC•ATP complexes maintain loaded Cy5-PCNA on all 5′ddPCy3/T•RPA complexes prior to the addition of Rad51•ATP complexes.

### Supplementary Methods

*Steady state FRET assays to monitor DNA-binding preference of Rad51*. A solution containing FRET P/T DNA (200 nM, **Figure S1**) and ATP was titrated with increasing concentrations of Rad51•ATP and E_FRET_ at each concentration was calculated. Specifically, at each Rad51•ATP concentration, *I*_665_ and *I_563_* are monitored over time until both stabilize for at least 1 min. Within this stable region, E_FRET_ values are calculated from the observed *I*_665_ and *I_563_* values and averaged to obtain the E_FRET_ value observed for a given concentration of Rad51•ATP.

*Pre-steady state FRET assays to monitor PCNA loading and unloading*. A solution containing 5′ddPCy3/T DNA (20 nM, **Figure S1**), neutravidin (80 nM homotetramer), and ATP is pre-incubated with RPA (25 nM heterotrimer). Then, Cy5-PCNA (20 nM homotrimer) is added, the resultant solution is transferred to a fluorometer cell, and the cell is placed in the instrument. *I*_665_ and *I_563_* are monitored over time until both stabilize for at least 1 min. Within this stable region, E_FRET_ values are calculated from the observed *I*_665_ and *I_563_* values and averaged to obtain the E_FRET_ value observed prior to addition of RFC•ATP complexes. Next, pre-formed RFC•ATP (20 nM RFC heteropentamer, 1 mM ATP) is added, the resultant solution is mixed via pipetting, and *I*_665_ and *I_563_* are monitored beginning ≤ 10 s after the addition of RFC•ATP (dead time ≤ 10 s) and continue to be until both stabilize for at least 1 min. Within this stable region, E_FRET_ values are calculated from the observed *I*_665_ and *I_563_* values and averaged to obtain the E_FRET_ value observed prior to addition of unlabeled PCNA. Finally, unlabeled PCNA (1.4 μM homotrimer) is added, the resultant solution is mixed by pipetting, and *I*_665_ and *I_563_* are monitored beginning ≤ 10 s after the addition of unlabeled PCNA (dead time ≤ 10 s).

### Supplemental Figures

**Figure S1.**
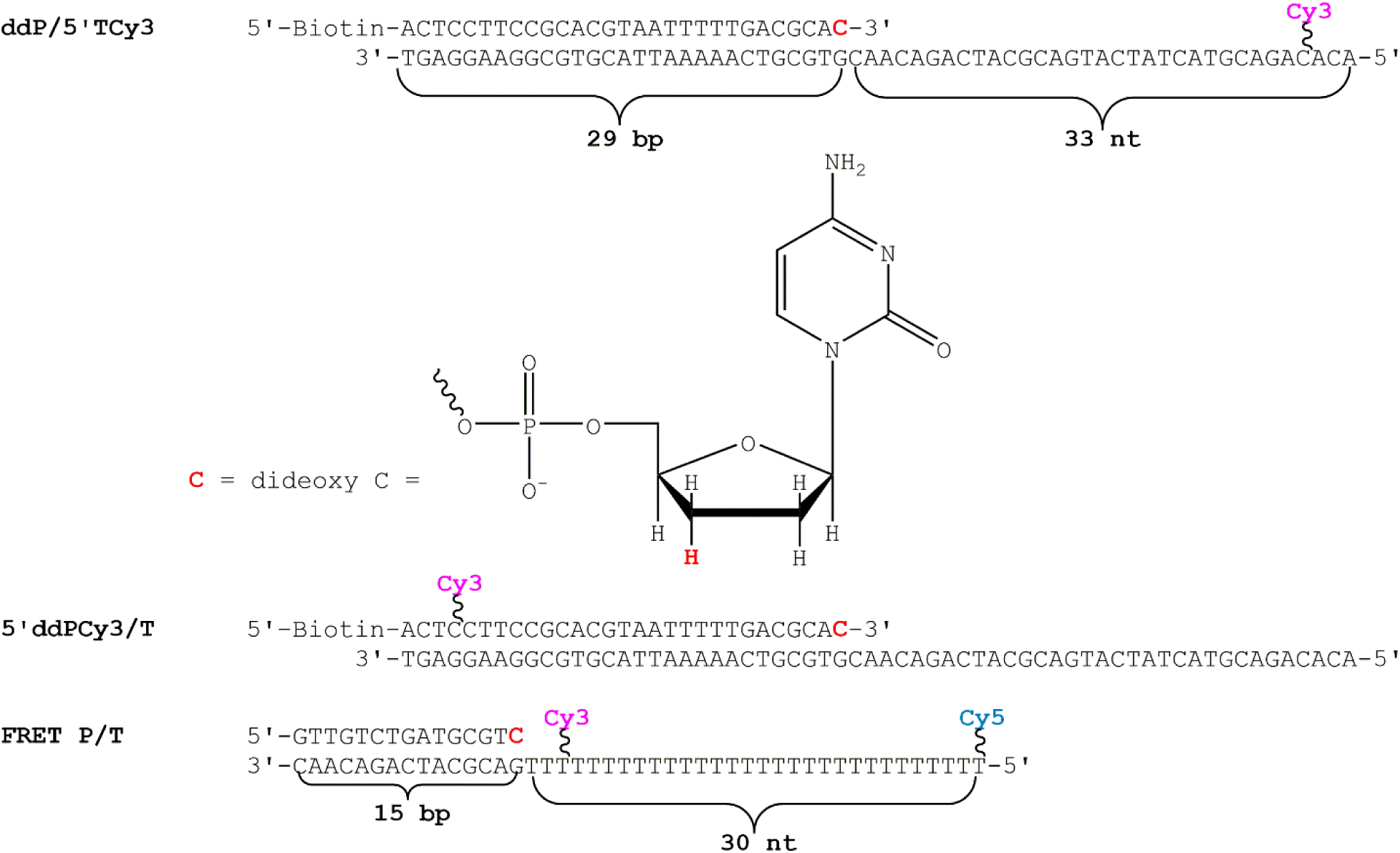
P/T DNA substrates utilized in this study. The sequences and lengths of the dsDNA and ssDNA regions are indicated. For the ddP/5′TCy3 and 5′ddPCy3/T DNA substrates, the size of the dsDNA region (29 bp) is in agreement with the requirements for assembly of a PCNA ring onto DNA by RFC^8,11,15^. The ssDNA regions of these substrates accommodate 1 RPA heterotrimer or 11 Rad51 monomers^1,2,16–18^. RPA prevents loaded PCNA from sliding off the ssDNA end of these substrates^8^. When pre-bound to neutravidin, the biotin attached to the 5′-end of the primer strands of these substrates prevents loaded PCNA from sliding off the dsDNA ends. Primers terminated at the 3′ end with a dideoxy C nucleotide cannot be extended by pol δ. For the FRET P/T DNA substrate, the dsDNA (15 bp) and ssDNA (30 nt) regions accommodate 5 and 10 Rad51•ATP, respectively^1,2^.

**Figure S2.**
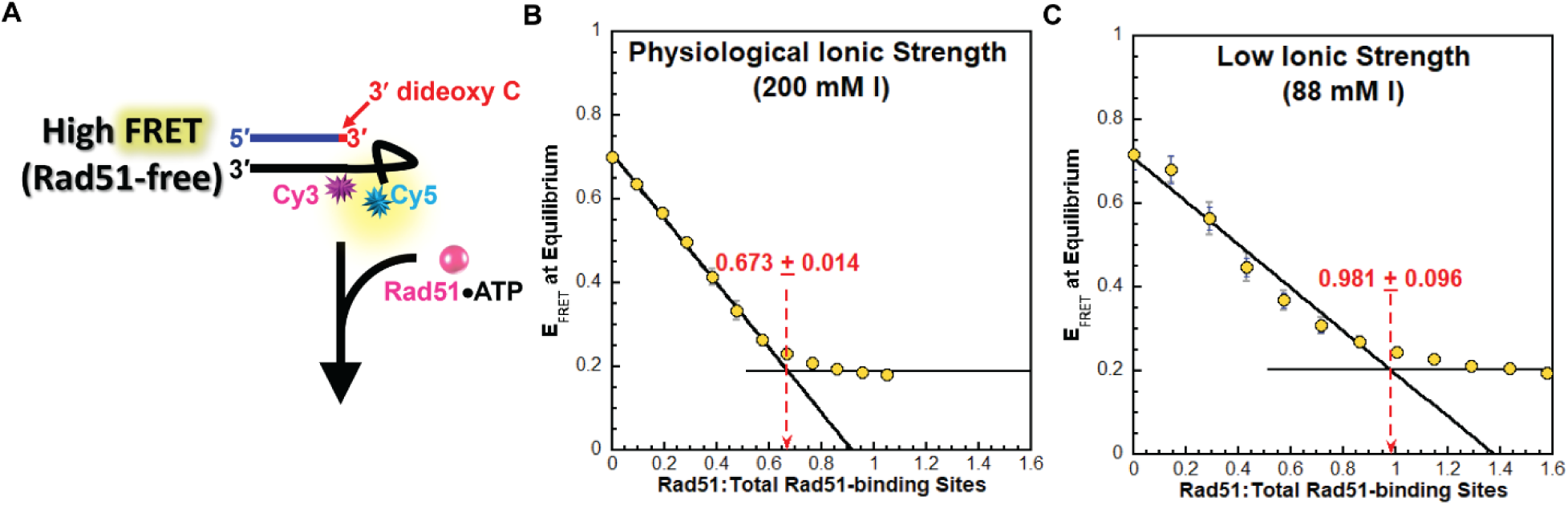
Rad51 has a strong, intrinsic preference for forming filaments on ssDNA over dsDNA at physiological ionic strength (**A**) Schematic representation of the FRET experiment utilizing a FRET P/T DNA substrate (**Figure S1**) and Rad51•ATP. The 30 nt ssDNA template (**Figure S1**) is labeled with an internal 3′ Cy3 (FRET donor) 3 nt from the P/T junction and a Cy5 (FRET acceptor) 3 nt from the 5′ terminus. In the absence of Rad51, the ssDNA template forms a collapsed, flexible structure, bringing the two cyanine fluorophores close together and yielding a high E_FRET_. Saturation of the ssDNA template with Rad51•ATP extends the engaged sequence ~1.6-fold relative to the B-form^7^, thereby increasing the Cy3–Cy5 distance and reducing E_FRET_. (**B** - **C**) FRET data. 200 nM FRET P/T DNA is titrated with Rad51•ATP at either physiological (200 mM I) or low ionic strength (88 mM I) and E_FRET_ values are monitored. The observed E_FRET_ values are plotted (on the y-axes) as functions of the ratios of active Rad51 to Total Rad51-binding sites (on the x-axes). Each data point represents the mean ± S.E.M. of three independent measurements. Data for each titration is fit to two segments lines; a linear regression with a negative slope and a flat line. The equivalence point (indicated with standard errors of the calculation) for each titration is calculated from the intersection of the two segment lines. Titrations carried out at physiological (200 mM I) and low ionic strength (88 mM) are displayed in panels **B** and **C**, respectively.

**Figure S3.**
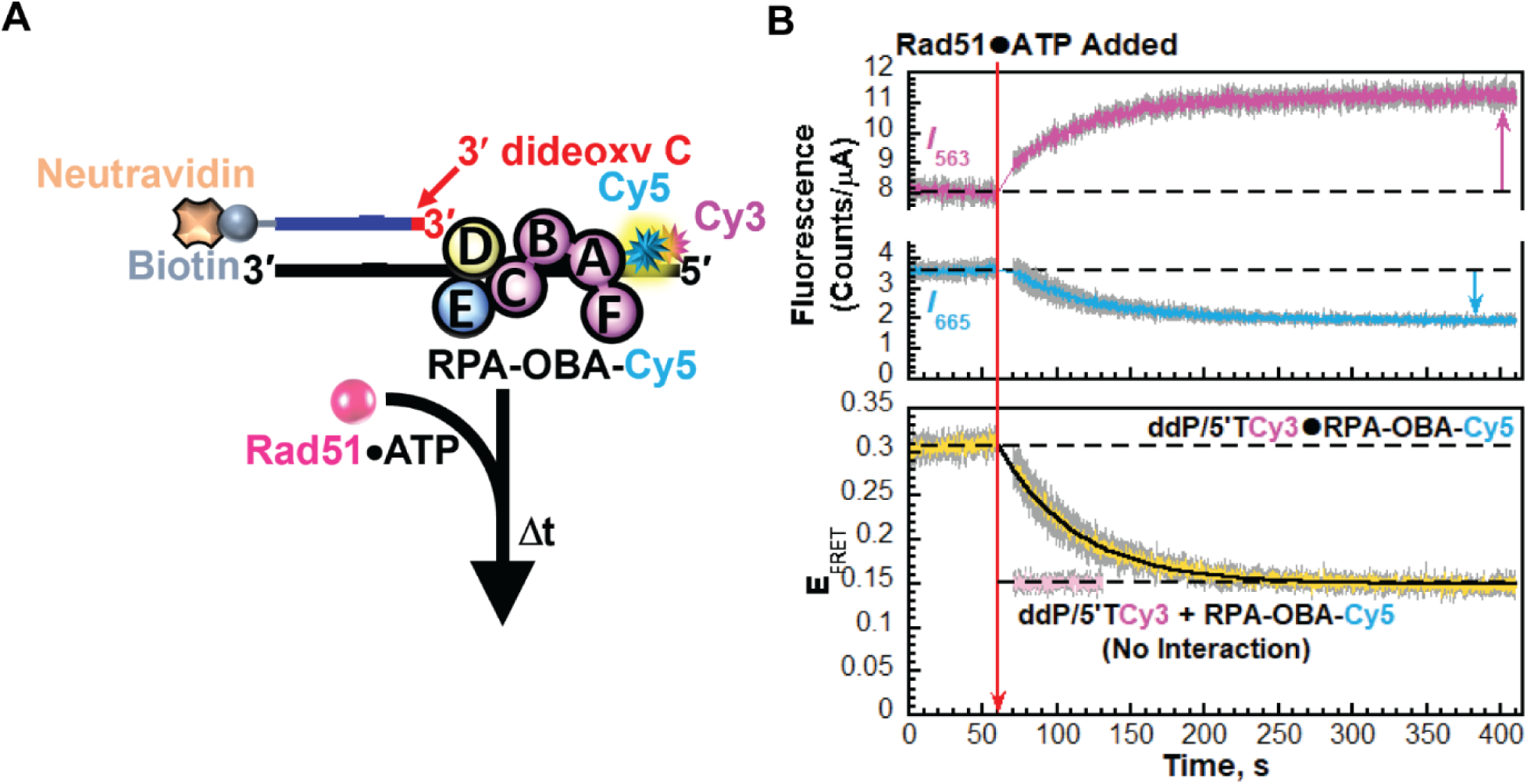
RPA/Rad51 exchange on P/T junctions devoid of PCNA. (**A**) Schematic representation of the FRET pair and experiment to monitor RPA/Rad51 exchange on P/T junctions devoid of PCNA. Experiments are performed exactly as described in the left panel of Figure 2A in the main text except RFC and PCNA were omitted. (**B**) Data. Each trace is the mean of at least three independent traces with the S.E.M. shown in grey. The time trajectories of *I*_563_ (magenta) and *I*_665_ (cyan) are displayed in the top panel and the corresponding E_FRET_ (yellow) is displayed in the bottom panel. The time at which Rad51•ATP is added is indicated by a red arrow. Changes in *I*_563_ and *I*_665_ are indicated in the top panel by magenta and cyan arrows, respectively. For observation, the *I*_563_ and *I*_665_, and the corresponding E_FRET_ values observed prior to the addition of Rad51•ATP complexes are each fit to dashed flat lines that are extrapolated to the axis limits. The predicted E_FRET_ trace (pink) for no interaction between RPA-OBA-Cy5 and the ddP/5′TCy3 DNA is displayed in the bottom panel, offset to the time after Rad51•ATP is added, and fit to a dashed flat line that is extrapolated to the axis limits. The rate constant for the E_FRET_ decrease (*k*_obs dec_) observed after the addition of Rad51•ATP complexes is reported in **Table 1** in the main text.

**Figure S4.**
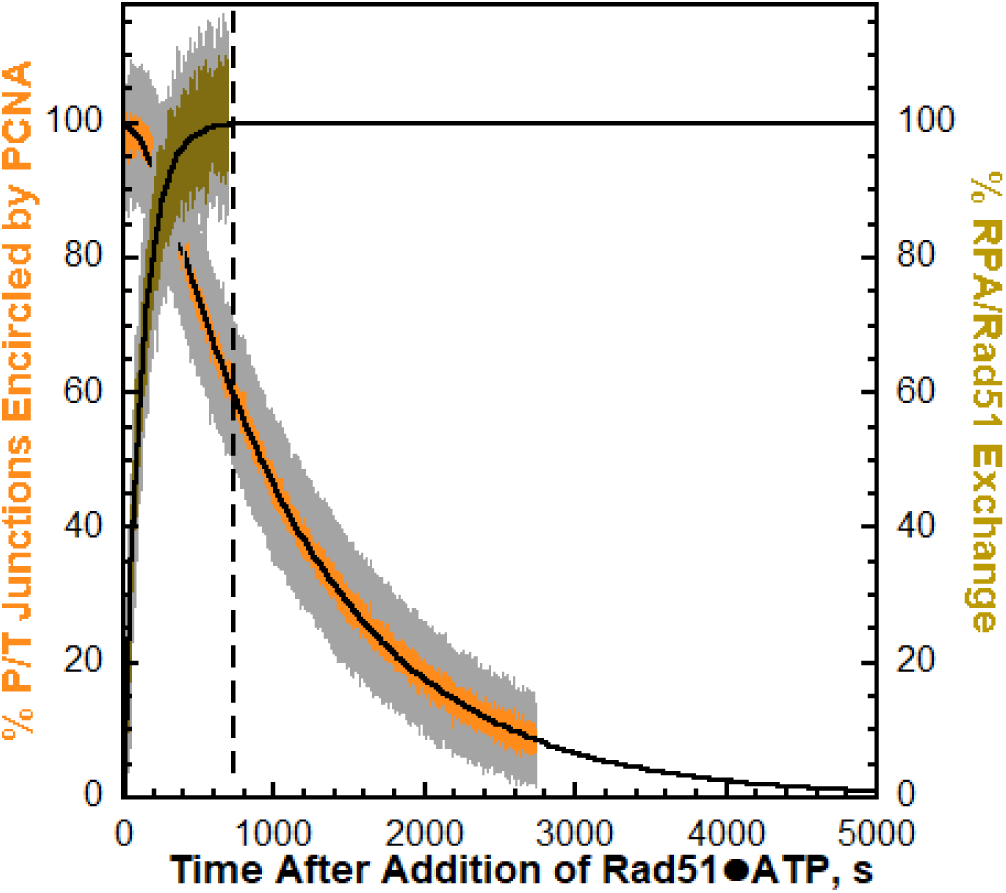
PCNA unloads from P/T junctions during RPA/Rad51 exchange and after RPA/Rad51 exchange has been completed. E_FRET_ traces from Figures 2B (RPA/Rad51 exchange) and **2C** (PCNA unloading) in the main text were each normalized to their respective ranges and the results plotted as a function of time (after Rad51•ATP is added) with the S.E.M. shown in grey. The normalized trace for RPA/Rad51 exchange (shown in brown) is fit to one-step, irreversible kinetic model that is extrapolated to the axis limits. The normalized trace for PCNA unloading (shown in orange), i.e. “% P/T Junctions Encircled by PCNA,” is fit to a two-step, irreversible kinetic model that is extrapolated to the axis limits. An extended time course is shown to highlight the extrapolated kinetics fits and lifetimes for complete unloading of PCNA from P/T junctions and complete RPA/Rad51 exchange on P/T junctions. For observation, the time at which RPA/Rad51 exchange is completed is indicated by a vertical dashed line.

**Figure S5.**
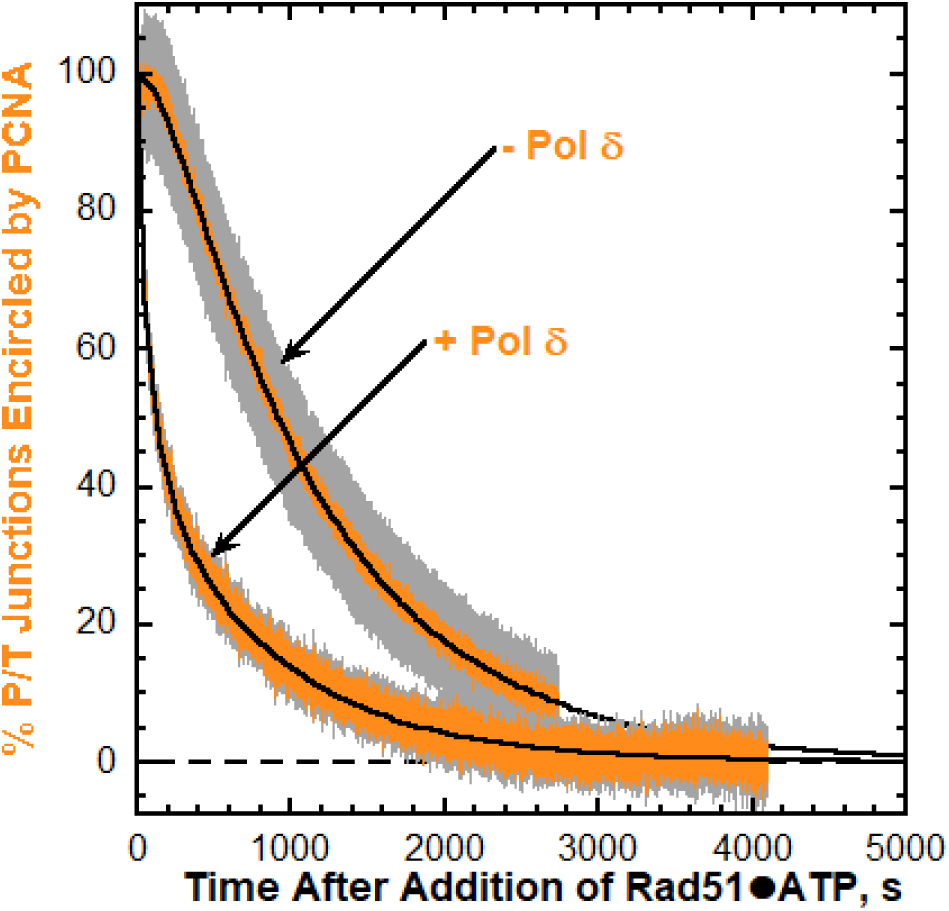
Pol δ holoenzymes do not stabilize loaded PCNA on P/T junctions during RPA/Rad51 exchange. E_FRET_ traces for unloading of PCNA from P/T junctions in the absence (Figure 2C) and presence of pol δ (Figure 3C) in the main text were each normalized to their respective ranges and the results plotted as a function of time (after Rad51•ATP complexes are added) with the S.E.M. shown in grey. The normalized traces for PCNA unloading, i.e., “% P/T Junctions Encircled by PCNA,” in the absence (-pol δ) and presence of pol δ (+ pol δ) are fit to a two-step, irreversible kinetic model and a triple exponential, irreversible kinetic model, respectively. Each kinetic model is extrapolated to the axis limits. A dashed flat line denotes 0% P/T junctions encircled by PCNA.

**Figure S6.**
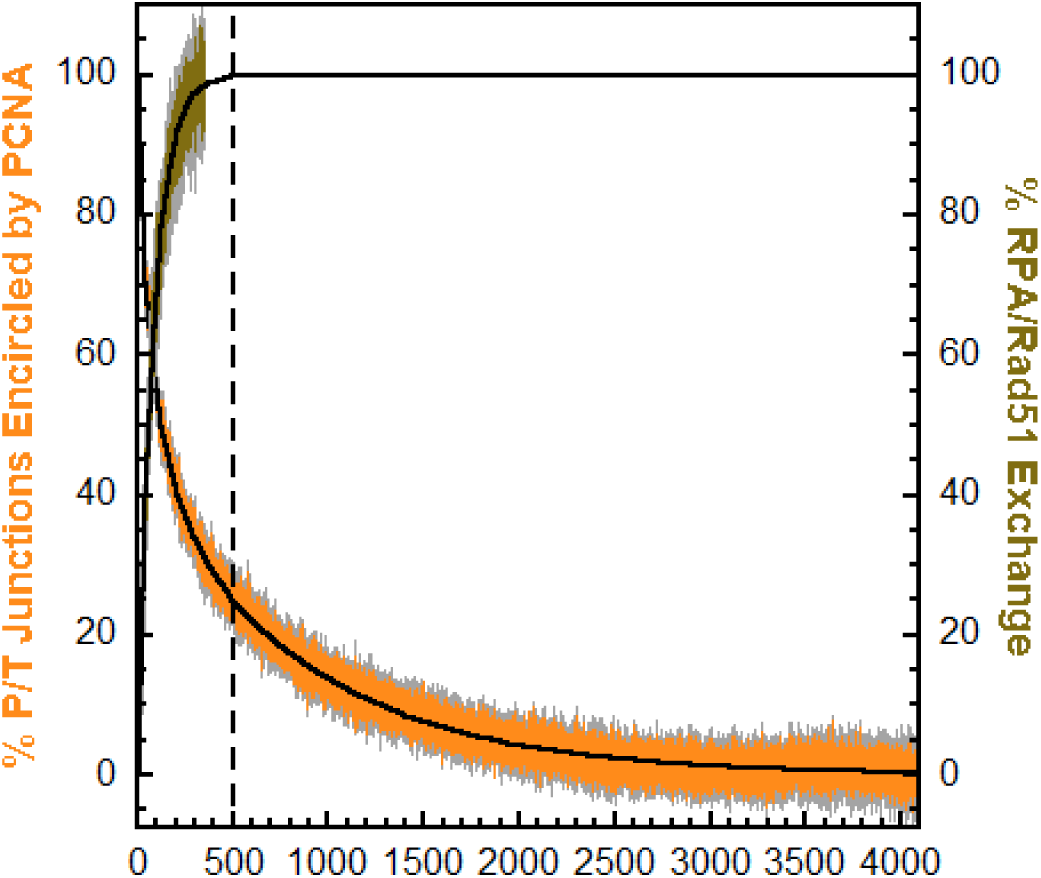
PCNA pre-engaged by pol δ in holoenzymes unloads from P/T junctions during RPA/Rad51 exchange and after RPA/Rad51 exchange has been completed. E_FRET_ traces from Figures 3B (RPA/Rad51 exchange) and **3C** (PCNA Unloading) in the main text were each normalized to their respective ranges and the results plotted as function of time (after Rad51•ATP complexes are added) with the S.E.M. shown in grey. The normalized trace for RPA/Rad51 exchange (shown in brown) is fit to a one-step, irreversible kinetic model that is extrapolated to the axis limits. The normalized trace for PCNA unloading (shown in orange), i.e., “% P/T Junctions Encircled by PCNA,” is fit to a triple exponential decline that is extrapolated to the axis limits. An extended time course is shown to highlight the extrapolated kinetics fits and lifetimes for complete unloading of PCNA from P/T junctions and complete RPA/Rad51 exchange on P/T junctions. For observation, the time at which RPA/Rad51 exchange is completed is indicated by a vertical dashed line.

**Figure S7.**
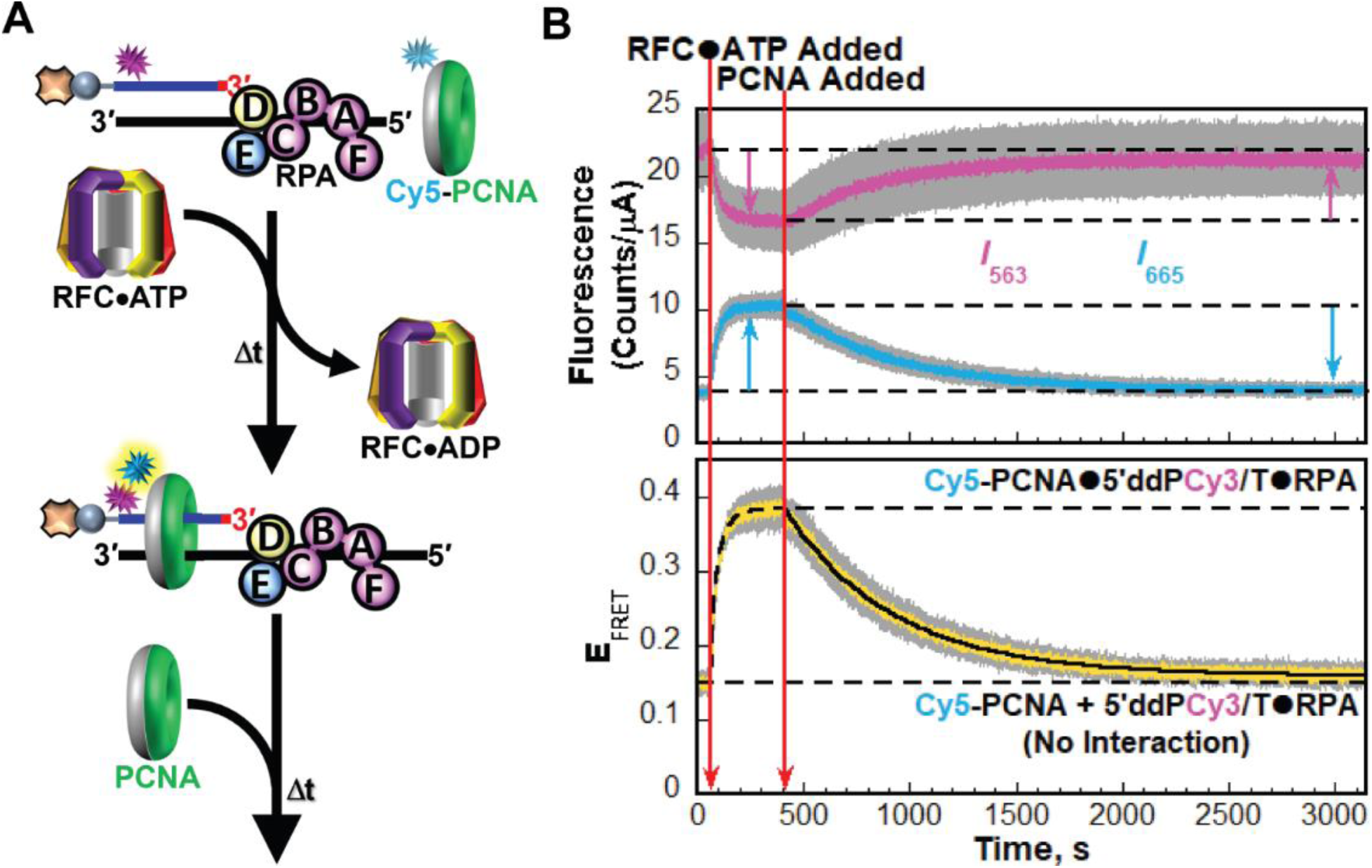
Dynamics of PCNA encircling P/T junctions engaged by RPA. (**A**) Schematic representation of the FRET pair and experiment to monitor the dynamics of PCNA encircling P/T junctions that are engaged by RPA. (**B**) Data. Each trace is the mean of at least three independent traces with the S.E.M. shown in grey. The time trajectories of *I*_563_ (magenta) and *I*_665_ (cyan) are displayed in the top panel and the corresponding E_FRET_ (yellow) is displayed in the bottom panel. The times at which RFC•ATP and unlabeled PCNA are added are indicated by red arrows. Changes in *I*_563_ and *I*_665_ are indicated in the top panel by magenta and cyan arrows, respectively. For observation, the *I*_563_ and *I*_665_, and the corresponding E_FRET_ values observed prior to the addition RFC•ATP are each fit to dashed flat lines that are extrapolated to the axis limits. Also, dashed flat lines are drawn to highlight the plateau *I* values observed after the addition of RFC•ATP. The increase in E_FRET_ observed upon the addition of RFC•ATP complexes is fit to a double exponential rise, yielding rate constants *k*_obs inc 1_ = 4.46 ± 0.2 (× 10^−2^) s^−1^ and *k*_obs inc 2_ = 1.70 ± 0.12 (× 10^−2^) s^−1^. The decrease in E_FRET_ observed upon the addition of unlabeled PCNA is fit to a single exponential decline, yielding the rate constant *k*_dec,obs_ = 1.93 ± 0.01 (× 10^−3^) s^−1^. The data for the PCNA loading portion of the plots is identical to that shown in Figure 4A in the main text.

## Footnotes

1 The content is solely the responsibility of the authors and does not necessarily represent the official views of the National Institutes of Health

## Notes

### Competing Interest Statement

The authors have declared no competing interest.

### Summary of Updates

Author order updated to clarify 1st author and corresponding author

